# AA147 Alleviates Symptoms in a Mouse Model of Multiple Sclerosis by Reducing Oligodendrocyte Loss

**DOI:** 10.1101/2025.01.01.630914

**Authors:** Metin Aksu, Kevin Kaschke, Joseph R. Podojil, MingYi Chiang, Ian Steckler, Kody Bruce, Andrew C. Cogswell, Gwen Schulz, Jeffery Kelly, R. Luke Wiseman, Stephen Miller, Brian Popko, Yanan Chen

## Abstract

Inflammation induced oligodendrocyte death and CNS demyelination, are key features of multiple sclerosis (MS). Inflammation-triggered endoplasmic reticulum (ER) stress and oxidative stress promote tissue damage in MS and in its preclinical animal model, experimental autoimmune encephalitis (EAE). Compound AA147 is a potent activator of the ATF6 signaling arm of the unfolded protein response (UPR) that can also induce antioxidant signaling through activation of the NRF2 pathway in neuronal cells. Previous work showed that AA147 protects multiple tissues against ischemia/reperfusion damage through ATF6 and/or NRF2 activation; however, its therapeutic potential in neuroinflammatory disorders remains unexplored. Here, we demonstrate that AA147 ameliorated the clinical symptoms of EAE and reduced ER stress, oligodendrocyte loss, and demyelination. Additionally, AA147 suppressed T cells in the CNS without altering the peripheral immune response. Importantly, AA147 significantly increased the expressions of *Grp78*, an ATF6 target gene, in oligodendrocytes, while enhancing levels of *Grp78* as well as *Ho-1*, an NRF2 target gene, in microglia. In cultured oligodendrocytes, AA147 promoted nuclear translocation of ATF6, but not NRF2. Intriguingly, AA147 altered the microglia activation profile, possibly by triggering the NRF2 pathway. AA147 was not therapeutically beneficial during the acute EAE stage in mice lacking ATF6 in oligodendrocytes, indicating that protection primarily involves ATF6 activation in these cells. Overall, our results suggest AA147 as a potential therapeutic opportunity for MS by promoting oligodendrocyte survival and regulating microglia status through distinct mechanisms.

## INTRODUCTION

Multiple sclerosis (MS) is an autoimmune demyelinating disease with diverse symptoms, often starting in early adulthood (Frohman et al., 2006; Reich et al., 2018). While the etiology of MS is currently unknown, recent studies indicate that oligodendrocyte apoptosis and axonal degeneration are key attributes of MS progression (Frohman et al., 2006; Lassmann, 2014). Current immunomodulatory therapies reduce relapse severity, but have limited impact on disease progression (Dargahi et al., 2017; Tintore et al., 2019). Complementary strategies are therefore urgently needed to protect the central nervous system (CNS) to slow down MS progression (Rodgers et al., 2013; Way and Popko, 2016; Titus et al., 2020).

Inflammation-induced endoplasmic reticulum (ER) stress and oxidative stress are thought to promote tissue damage in MS (Pegoretti et al., 2020). Inflammatory factors, such as tumor necrosis factor alpha (TNF-α) and interferon-gamma (IFN-γ), trigger ER stress in the CNS and activate the innate unfolded protein response (UPR) (Cunnea et al., 2011; Stone and Lin, 2015). The UPR is regulated by activating transcription factor 6 alpha (herein referred to as ATF6), protein kinase RNA-like endoplasmic reticulum kinase (PERK), and inositol-requiring enzyme 1 (IRE1), and is responsible for maintaining cellular homeostasis following acute ER insults. However, prolonged ER stress can lead to apoptosis (Walter and Ron, 2011; Hetz et al., 2020). In addition to ER stress, oligodendrocyte and myelin damage can also be caused by oxidative stress, which is amplified by activated microglia and astrocytes (Haider et al., 2011; Mendiola et al., 2020; Steudler et al., 2022). The endogenous antioxidant response system protects cells from oxidative stress by increasing the expression of cytoprotective enzymes primarily through the activity of the nuclear factor erythroid 2– related factor 2 (NRF2) (Motohashi and Yamamoto, 2004).

Among the three arms of the UPR, the mechanism behind ER stress-dependent ATF6 activation is the least understood. At rest, ATF6 mainly forms disulfide-bound dimers/oligomers, maintained by ER lumen-localized protein disulfide isomerases (PDIs). Under ER stress, PDIs reduce these disulfide bonds, leading to ATF6 oligomer dissociation, and subsequent trafficking to the Golgi, where it undergoes sequential cleavage by site 1 and site 2 proteases (S1P and S2P) (Ye et al., 2000). This releases the cytosolic domain of ATF6, which localizes to the nucleus and enhances cellular protection against ER stress by upregulating the transcription of certain cytoprotective ER chaperones, such as GRP78 or BIP, ER associated degradation (ERAD) components, and antioxidant proteins (Adachi et al., 2008; Shoulders et al., 2013; Jin et al., 2017). Intriguingly, ATF6 is activated in MS patient lesions and MS-like animal models (Mháille et al., 2008; Stone and Lin, 2015). However, the precise contribution of ATF6 signaling pathway in MS-relevant cell types remains unclear.

Recently, a pharmacologic ATF6 activator, N-(2-hydroxy-5-methylphenyl)-3-phenylpropanamide (AA147), was discovered using a high-throughput screening approach (Plate et al., 2016; Paxman et al., 2018; Kline et al., 2023). Compound metabolic activation and subsequent covalent modification of a subset of PDIs by AA147 disrupts PDI-dependent maintenance of ATF6 disulfide bonds, leading to more reduced ATF6 monomers that can traffic to the Golgi for proteolytic activation (Paxman et al., 2018). AA147 provides protection against ischemia/reperfusion damage to the heart, liver, kidney, and brain, the latter indicating that this compound can cross the blood-brain barrier (Blackwood et al., 2019). Although AA147 was originally developed as a pharmacological activator of the ATF6 arm of the UPR, recent findings show that AA147 shields neuronal cells against glutamate-induced oxidative stress primarily through its activation of the NRF2 antioxidant response pathway in neural cell types (Rosarda et al., 2021; Kline et al., 2024). AA147 also provides neuroprotection post-cardiac arrest, likely through both ATF6 and NRF2 pathway activation (Yuan et al., 2022). NRF2 is normally bound in the cytosol to the Kelch-like ECH-associated protein 1 (KEAP1) that targets it for degradation. In neural cells, AA147 activates NRF2 by covalently modifying KEAP1, diminishing KEAP1-NRF2 binding and increasing NRF2 nuclear translocation (Rosarda et al., 2021). Nevertheless, the therapeutic potentials and underlying mechanisms of AA147 have not yet been explored in neuroinflammatory disorders.

In this study, we find that AA147 treatment significantly reduces the clinical symptoms of EAE, and the protection provided by this compound correlates with reduced oligodendrocyte loss and demyelination. We also show that AA147 suppresses CNS inflammation but does not alter peripheral immune cell activation and infiltration. Importantly, AA147 significantly increases the expressions of *Grp78*, an ATF6 target gene, in oligodendrocytes, while enhancing levels of both *Grp78* and the antioxidant heme oxygenase-1 (*Ho-1*), an NRF2 target gene, in microglia. Moreover, we observe the nuclear translocation of ATF6, but not NRF2, in cultured oligodendrocytes treated with AA147, indicating that AA147 selectively activates ATF6 in this cell type. Intriguingly, we demonstrate that AA147 reduces apoptotic cell death of oligodendrocytes and facilitates a microglial profile shift. Finally, we show that oligodendrocytic deletion of ATF6 mitigates AA147 protection early in disease pathogenesis, indicating that AA147-dependent activation of ATF6 in oligodendrocytes is a major contributor to the protection afforded by this compound. Collectively, our results define adaptations induced by AA147 in specific glial cell types and demonstrate the therapeutic potential for this compound in mitigating pathologies in the EAE mouse model of MS.

## MATERIALS AND METHODS

### Transgenic Animals and Tamoxifen Administration

*PLP-CreER^t^* mice (Doerflinger et al., 2003), were crossed with *ATF6^fl/fl^ floxed* mice to obtain a tamoxifen induced conditional knockout mouse line – *ATF6^fl/fl^; PLP-CreER^t^*. Tamoxifen was injected intraperitoneally prior to EAE induction at six to seven weeks of age for five consecutive days with a single daily IP injection of 1mg/day of tamoxifen (Hussien et al., 2014). After three weeks of recovery to clear tamoxifen, the mice were subjected to EAE induction at 10-week-old.

### EAE Induction

EAE was induced in around 10-week-old female C57BL/6J mice or transgenic mice by subcutaneous flank administration of 200 µg MOG_35-55_ peptide emulsified with complete Freund’s adjuvant (CFA) (MOG_35-55_/CFA), supplemented with inactive *Mycobacterium tuberculosis* H37Ra (Hooke Laboratories, Cat. # EK-2110). IP injections of 150 ng of pertussis toxin were given one hour after MOG emulsion administration and again, 24 hour later. Mice were weighed and scored daily for clinical signs as follows: 0 = healthy, 1 = flaccid tail, 2 = ataxia and/or paresis, 3 = paralysis of hindlimbs and/or paresis of forelimbs, 4 = tetraparalysis, 5 = moribund or death. Mouse groups were randomized before the treatment.

### Drug Administration and Tissue Collection

AA147 working solution was prepared in 10% DMSO, 10% Koliphor EL:EtOH, and 80% saline. To optimize dosage, 2mg/kg and 8mg/kg of AA147 were administered through the tail vein, gavage or IP. Lumbar spinal cords were freshly collected after 24 hours. For EAE treatment, the mice were injected intraperitoneally with AA147 or the equivalent amount of vehicle daily starting from PID 7, and clinical symptoms were recorded daily. Lumbar spinal cords were collected from a separate cohort at PID 12 or PID 16.

### Flow Cytometry

The spleens and lymph nodes were processed via physical disruption, and and for spleen RBCs were lysed using ammonium chloride prior to processing for flow cytometry. For the CNS, single cell suspensions were prepared by mincing the tissue in 2 mL of Accutase (MilliPore), and the samples were incubated at 37°C for 30 min. Single cells were separated from the myelin and cellular debris via centrifugation in a 40% percoll solution at 650xg for 25 minutes. Following the enzyme digestion, the CNS samples were disrupted with a 100 μm cell strainer, which was then twice washed with 10 mL of HBSS+ 5% FCS, followed by pelleting the cells. For intracellular staining, cells (1×10^6^ cells in 200 µL of complete RPMI medium) were cultured in the presence of PMA (50 ng/mL), ionomycin (500 ng/mL) for two hours, and brefeldin A (10 µg/mL) for an additional two hours. Cells were collected and a live/dead stain was performed per the manufacturer’s instructions (Invitrogen, L34955). Following live/dead staining, the cells were incubated in flow cytometry staining buffer (1x PBS + 5% FCS) plus Fc Block (Anti-Mouse CD16/CD32 Purified; eBioscience) in the dark at 4°C for 30 minutes. Samples were washed three times in 1x PBS + 5% FCS, and then the samples were incubated with antibodies for cell surface staining (diluted 1:200 in 1x PBS + 5% FCS) in the dark at 4°C for 30 minutes. For intracellular staining, a fixation/permeabilization (Intracellular Fixation & Permeabilization Buffer Set; ThermoFisher) working solution was added to each tube and pulse vortexed. Samples were incubated in the dark at 4°C for 60 minutes. Samples were washed three times in 1X permeabilization buffer volume after decanting and were blocked with 2% normal rat serum by adding 2 µL directly to the cells. The samples were then incubated at room temperature for 15 minutes. Without washing, the fluorochrome-conjugated antibodies (diluted 1:200 in 1x permeabilization buffer) for detection of intracellular antigen(s) were added to the cells and samples were incubated in the dark at room temperature for 30 minutes. Samples were then washed three times in 1X permeabilization buffer, followed by three washes with 1x PBS. Stained cells were resuspended in 1x PBS + 5% FCS, and samples were run on a BD Celesta (BD Bioscience), and the data analyzed using FloJo Version 10.10.0 software (Tree Star, Inc., Ashland, OR). The specific antibodies used are presented in Supplementary Table S1.

### Ex Vivo Recall

For *ex vivo* recall cultures, two sets of cultures were set up, with cells isolated as described above from the inguinal lymph nodes and spleens. Cells were cultured in triplicate wells per individual mouse (*n*=5 mice per treatment group) at 5 × 10^5^ cells per well in the presence of anti-CD3 (1 μg/ml), OVA_323–339_ or MOG_35–55_ (20 μg/ml) in the complete RPMI medium. For cytokine analysis, replicate wells were harvested on day+3 of culture and the level of cytokine secreted determined via multiplex Luminex Liquid Chip (Millipore).

### Adoptive Transfer EAE Immunization

For the adoptive transfer EAE experiments, draining lymph nodes from MOG_35-55_/CFA primed C57BL/6J mice as described above were collected on PID8. Isolated cells from lymph nodes were reactivated in the presence of MOG_35–55_ (20 μg/ml) and IL-12 (10 ng/ml) in complete RPMI medium. After 72 h in culture, total cells were counted labeled with carboxyfluorescein succinimidyl ester (CFSE), the number of total number cell versus blast cells were re-counted, with blast cells typically representing 30% of total cell numbers. The cells were resuspended in 1xPBS and injected intravenously into the tail vein of 8-10-week-old naïve C57BL/6J female mice, with each recipient receiving 3 × 10⁶ blast cells. Recipient mice were given two IP injections of 200 ng pertussis toxin each 0 and 48 h later and randomly assigned to either vehicle or AA147 (8mg/kg) treatment. After 6 days of treatment, spleen and CNS, include brain and spinal cord, were then collected for flow cytometry analysis. Transferred T cells was determined by CFSE dilution.

### Immunofluorescent Staining

Four percent paraformaldehyde fixed frozen sections were cut in a cryostat at 10 μm thickness. Sections were allowed to dry and were permeabilized with 1x PBS-T (0.2% Triton-X-100) for 20mins at RT. Sections were blocked with a blocking buffer (1% donkey serum, 5% BSA, 2% fish gelatin, 0.1% Triton-X-100) for one hour at RT and incubated with primary antibodies overnight at 4°C. Primary antibodies were washed off with 1X PBS, and Alexa fluor-conjugated secondary antibodies (Thermo Fisher Scientific) were allowed to incubate on sections for 1 hour at RT. After 1xPBS washing, sections were mounted in Fluoromount-G with DAPI (Thermo Fisher Scientific, 00495952). The primary antibodies used in this study are as follows: iNOS (1:100, BD Lab, 610328), MBP (1:100, Bio-Rad, MCA409S), Cleaved Caspase 3 (1:400, Cell Signaling, 9661T), CoraLite Plus 488 Anti Mouse CD11b (1:300, CoraLite Plus, CL488-65055), TPPP (1:50, Invitrogen, PA5-19243), NF-H (1:500, Millipore, AB5539), GFAP (1:200, Millipore, AB5541), Sox 10 (1:50, R&D, AF2864), ATF-6a (1:50, Santa Cruz Biotechnology, sc-166659), Arg1 (1:50, Santa Cruz Biotechnology, sc-166920), CD3 (1:50, Santa Cruz Biotechnology, sc-20047), GRP 78 (1:50, Santa Cruz Biotechnology, sc-166490), HO-1 (1:50, Santa Cruz Biotechnology, sc-390991), Nrf2 (1:50, Santa Cruz Biotechnology, sc-365949), and TMEM 119 (1:250, Synaptic Systems, 400 004). The slides were imaged with an Olympus Slideview VS200 slide scanner and/or with a Zeiss LSM 880 Confocal system. Mean fluorescent indices were calculated using QuPath software with a threshold-based approach (Bankhead et al., 2017). Appropriate thresholds for cell markers were determined and used to find pixel coordinates that were localized within the cells of interest. Fluorescent intensities were then taken from the localized pixel coordinates and used to find the average intensity values of corresponding protein immunofluorescence. Additionally, Qupath software was used to count cells using the Positive Cell Detection tool. Appropriate thresholds for nuclei detection, cytosolic/nuclear compartmental thresholds, and cell expansion were determined for different cell types and stains.

### In Situ Hybridization

Fixed frozen sections were allowed to dry. An ACD RNAScope Multiplex V2 protocol (ACDBio) was followed for triplex fluorescent ISH labeling. ACD probes Myrf-C1, Hmox1-C2 and Grp78-C3 (ACDBio) were used to hybridize target mRNAs. TSA VIVID Dyes 520, 570, 650 were used in combination (Tocris Bioscience). Slides were imaged on a Zeiss LSM 880 Confocal System. Images were analyzed to find approximate dot counts in Qupath using the subcellular detection method. Appropriate thresholds for nuclei detection, subcellular dots, and cell expansion were determined for the project and used to automatically measure approximate dot quantities. Data was purged of all cells with no or little Myrf expression (<3 dots). Calculated dots were then averaged over a single sample (>30 cells per sample).

### DHE Assay

Fixed frozen sections were allowed to dry. Slides were then rinsed to wash out the OCT compound. Slides were immediately placed into 5 µM DHE solution (Sigma, D7008) and incubated at 40°C for one hour. After ddH2O washing, sections were mounted in Fluoromount-G (Thermo Fisher Scientific, 00495802).

### Oligodendrocyte Precursor cell (OPC) Culture

Isolation and immunopanning purification were performed as previously described (Dugas and Emery, 2013). Briefly, mouse brain cortices from seven-day-old pups were diced and digested with papain at 37°C. Cells were triturated and resuspended in a panning buffer, and then incubated at room temperature sequentially on plates coated with primary antibodies against rat neural antigen-2 (Ran-2) and galactosylceramide (GalC) for negative selection and O4 for positive selection. OPCs were released from the final panning dish with trypsin and seeded on poly-D-lysine-coated flasks in growth medium to proliferate. OPCs were then split and plated in differentiation media with T3 overnight. Differentiating cells were treated with AA147 (10 μM) and/or recombinant IFN-γ (200 U/ml, R&D systems) for 16 hours before fixation.

### Immunocytochemistry

Four percent paraformaldehyde culture slides were permeabilized in 1xPBS-T (0.2% Triton-X-100) and then blocked with blocking buffer (1% donkey serum, 5% BSA, 2% fish gelatin, 0.1% Triton-X-100) for 30mins at RT. Primary antibodies were applied overnight at 4°C and followed by incubation with Alexa fluor-conjugated secondary antibodies for two hours at RT. Sections were mounted in Fluoromount-G with DAPI and allowed to dry overnight at 4°C prior to imaging on a Zeiss LSM 880 Confocal System.

### RT-quantitative PCR (RT-qPCR)

The RT-qPCR procedure was described previously (Chen et al., 2019). Total RNA was isolated from the lumbar spinal cords with Aurum™ Total RNA Fatty and Fibrous Tissue Kit (Bio-Rad, 7326830) and then reversed into cDNA with an iScript™ cDNA synthesis kit (Bio-Rad,1708891). RT-qPCR was performed on a Thermo QuantStudio 3 (Thermo Fisher Scientific) using SYBR Green technology. Results were analyzed and presented as the fold-induction relative to the internal control primer for the housekeeping gene RPL13A. All primers used in this study are listed in Supplementary Table S2.

### Xbp1 Splicing Assay

RNA extraction and cDNA synthesis were performed as described in RT-qPCR. Xbp1 cDNA was amplified with DNA polymerase using primers across the splice site capable of amplifying both unspliced and spliced forms of xbp1, separated by electrophoresis on a 2% agarose gel. The size difference between the spliced and the unspliced XBP1 is 26 nucleotides. XBP1 primers were used as described (He et al., 2017): forward primer, CAGAGTAGCAGCGCAGACTGC; and reverse primer, TCCTTCTGGGTAGACCTCTGGGAG. The amplification products were separated by DNA electrophoresis. 3T3 cell lines treated with ER stress inducer thapsigargin (200 nM, Sigma, T9033) were used as a positive control.

### Western-Blot

Fresh frozen spinal cord tissue was lysed in ice-cold RIPA buffer (Sigma) containing Halt protease and phosphatase inhibitor cocktail (Thermo Fisher Scientific, PI78441). Lysates were centrifugated at 12,000 RPM for 20min at 4°C. A total 17 µg of protein lysates was separated by 4– 12% SDS-PAGE (Bio-Rad, 4561095) and transferred to a nitrocellulose membrane. The membranes were blocked in 5% non-fat milk. Primary antibodies were incubated at 4C overnight and washed with 1x PBS-T. After incubating with HRP conjugated second antibodies for 1 h at RT and washing with 1x PBS-T, the membranes were developed with ECL reagents and scanned by the iBright CL1500 Imaging System (Thermo Fisher Scientific). The resulting bands were analyzed using iBright Analysis Software. The primary antibodies used for Western blot are as follows: Mouse anti-CHOP (1:500, Invitrogen, MA1-250), HRP beta actin mouse monoclonal antibody (1:10000, Proteintech, HRP-60008), rabbit anti-cleaved caspase 3 (1:400, Cell Signaling, 9661T), rabbit anti-BIP (1:1000, Cell Signaling, C50B12), mouse anti-HO-1 (1:500, Santa Cruz Biotechnology, sc-390991), and rabbit Bax (1:500, Cell Signaling, 2772T).

### Protein Carbonyl Assay

Tissue homogenates were derivatized by 2,4-dinitrophenylhydrazine (DNPH) using the Protein Carbonyl Assay Kit (Abcam, ab178020). Then, 7μg of derivatized lysate was separated by a 4-12% SDS-PAGE (Bio-Rad, 4561095) and transferred to a nitrocellulose membrane. The membrane was incubated with Ponceau before being blocked with 5% fat-free milk, followed by the application of anti-dinitrophenyl antibody and then HPR conjugated secondary antibodies provided in the kit. The development and imaging of membranes were carried out using the same Western blot protocol as described above.

### Statistical Analysis

All results are expressed as mean±SEM. Statistical significance for these data was determined by t-tests, analysis of variance (ANOVA) followed by Turkey’s posttest for multiple group experiments using Graphpad Prism 10. Significance levels were set at *p < 0.05, **p < 0.01, ***p < 0.001, ****p < 0.0001.

## RESULTS

### AA147 alleviates EAE symptoms and reduces loss of oligodendrocytes, myelin and axons

To determine the optimal AA147 dosing strategy in the CNS, we administrated 2mg/kg and 8mg/kg of AA147 to C57BL/6J female mice via gavage, intraperitoneal (IP) and intravenous injections (Figure 1A and Supplementary Figure 1A, B). Lumbar spinal cords were collected after 24 hours for RT-qPCR analysis. Our results showed that between two dosages, IP injections of AA147 at 8mg/kg, significantly induced the expression of ATF6 downstream targets *Grp78*, *Grp94*, and *Pdia6* (p<0.05, p<0.0001, and p<0.01 respectively), but not the PERK-regulated gene *Atf4* or the IRE1-regulated gene *Erdj4* (Figure 1A). Similarly, we observed a significant increase in the expression of *Nrf2* (p<0.05) as well as its downstream targets *Ho-1* (p<0.01) with the same dosing strategy (Figure 1B). However, no significant changes on the expression of these target genes were found in the spinal cords obtained from mice administrated AA147 by gavage or intravenous injections, except that 2mg/kg of AA147 by gavage increased the *Grp78* levels (p<0.05) (Supplementary Figure 1A, B).

**Figure 1:**
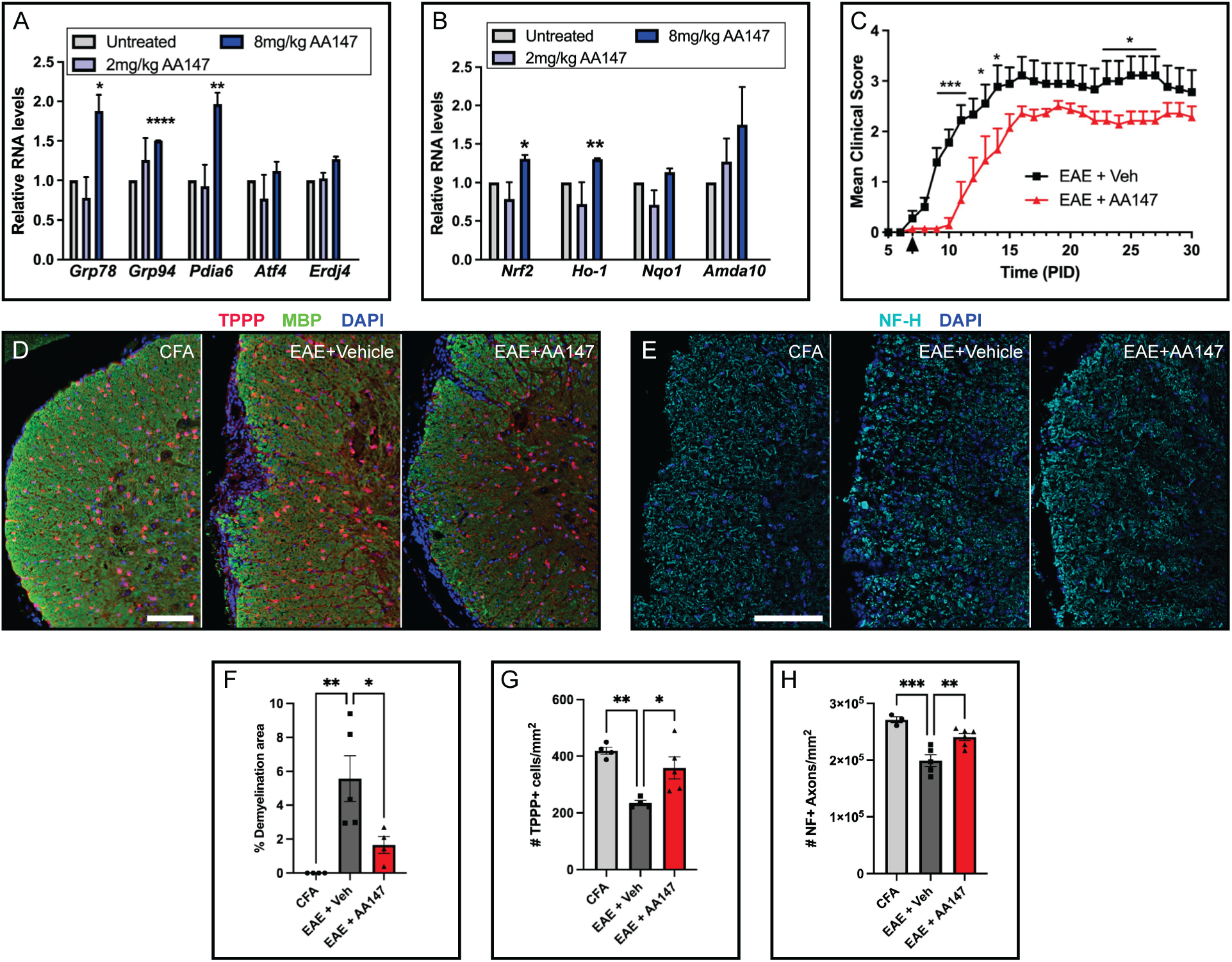
AA147 activates ATF6 and NRF2 pathways and ameliorates EAE. (A, B) Total RNA was isolated from lumbar spinal cord tissue of C57BL/6J female mice 24 hours after IP injections of AA147. (A) Relative mRNA expressions of UPR genes and (B) NRF2 downstream target genes were determined by RT-qPCR. Data are expressed as mean ± SEM; n= 3 per group; *p< 0.05, **p< 0.01, ****p<0.0001 by one-way ANOVA. (C) Clinical scores of C57BL/6J female mice immunized with MOG_35-55_/CFA to induce EAE, treated with IP of vehicle (n = 9) and 8mg/kg AA147 (n = 7). Treatments was started at PID 7 (arrow pointed). *p< 0.05, ***p< 0.001 by multiple t-tests. (D-H) Histology analysis of cross-sections of lumbar spinal cord collected at PID 16 from CFA mice or EAE mice treated daily with IP injection of vehicle or 8mg/kg AA147. (D) Representative images of TPPP and MBP staining (Scale bar=100µm), (F) quantification of the percentage of demyelinated area and (G) the number of TPPP+ oligodendrocytes in the white matter of the lumbar spinal cord. (E) Representative images of neurofilament (NF) staining (Scale bar=100µm) and (H) quantification of the number of NF+ axons in the white matter of lumbar spinal cord. Data are expressed as mean ± SEM; n= 4-5 per group; *p< 0.05, **p< 0.01, ***p<0.001 by one-way ANOVA.

Next, we used 8mg/kg (IP) to determine whether AA147 can improve the outcomes of EAE. C57BL/6J female mice immunized with MOG_35-55_/CFA underwent a 30-day testing period, during which we administered 8mg/kg of AA147 or vehicle (IP) to EAE mice daily starting from post-immunization day 7 (PID 7). The first clinical signs of EAE (clinical score 1.0) in mice treated with vehicle (EAE+Veh) appeared at PID 8, with the disease course peaking at PID 15. EAE animals treated with AA147 (EAE+AA147) showed significantly milder disease severity and reached peak clinical scores at PID 19 (Figure 1C).

To observe the pathological effects of AA147 treatment on the CNS, we used EAE animals from a separate treatment cohort, and isolated lumbar spinal cords at PID16. Western blot analysis revealed that EAE+AA147 mice showed significantly higher levels of GRP78/BIP than EAE+Veh mice, indicating activation of the ATF6 pathway (Supplementary Figure 2). Demyelinated lesions measured by MBP-absent white matter significantly increased in EAE+Veh mice (5.6%) compared to mice immunized with only adjuvant (CFA) (p<0.01), while AA147 treatment limited the demyelinated area to about 1.7% (p<0.05) (Figure 1D, F). Furthermore, we tested whether AA147 confers oligodendrocyte resistance to inflammatory attacks and prevents axon damage. Compared to CFA mice, EAE+Veh mice lost about 44% of TPPP+ mature oligodendrocytes (p<0.01) and 27% neurofilament+ axons (p<0.001) (Figure 1D, E, G, H). However, compared to vehicle, AA147 treatment prevented the loss of TPPP+ oligodendrocytes (p<0.05) and axons (p<0.01) in the white matter of lumbar spinal cords at PID16. We also examined the neuropathology changes at an earlier time point (PID12), and our results indicate that there were no significant differences between vehicle and AA147 treatment groups regarding the density of CD3+ T cells, NF+ axons, TPPP+ oligodendrocytes, or demyelinated lesion areas (Supplementary Figure 3). These findings indicate that AA147 ameliorates EAE severity, which correlates with oligodendrocyte survival, myelination, and axon integrity at the peak of disease.

### AA147 does not affect the peripheral immune response

Since AA147 treatment reduced EAE severity, we next sought to determine the underlying mechanism of this protection. EAE is mediated by myelin protein-specific T cells, which are activated in secondary lymphoid organs, such as lymph nodes and spleen. These T cells then translocate into the CNS through a compromised blood brain barrier, inducing CNS inflammation and demyelination (Furtado et al., 2008). To examine whether AA147 directly affects the activation of the peripheral immune response, we isolated cells from the spleen and lymph nodes of EAE mice at PID 12. Compared with vehicle-treated EAE mice, EAE mice treated daily with 8mg/kg of AA147 did not show a difference in the numbers of CD4+ T cells, B cells, CD11b+ cells, and antigen presenting cells (APCs) (macrophages, monocytes, neutrophils and dendritic cells) in the spleen as well as in lymph nodes as determined by flow cytometric analyses (Figure 2A, B). In addition, AA147 treatment had no effect on IFN-γ or interleukin (IL)-17 cytokine production of splenocytes (Figure 2C, E) and lymph nodes (Figure 2D, F) in response to anti-CD3, OVA_323–339_ or MOG_35–55_. Together, the above findings suggest that AA147 does not impact immune cell activation in the peripheral tissues of EAE mice.

**Figure 2:**
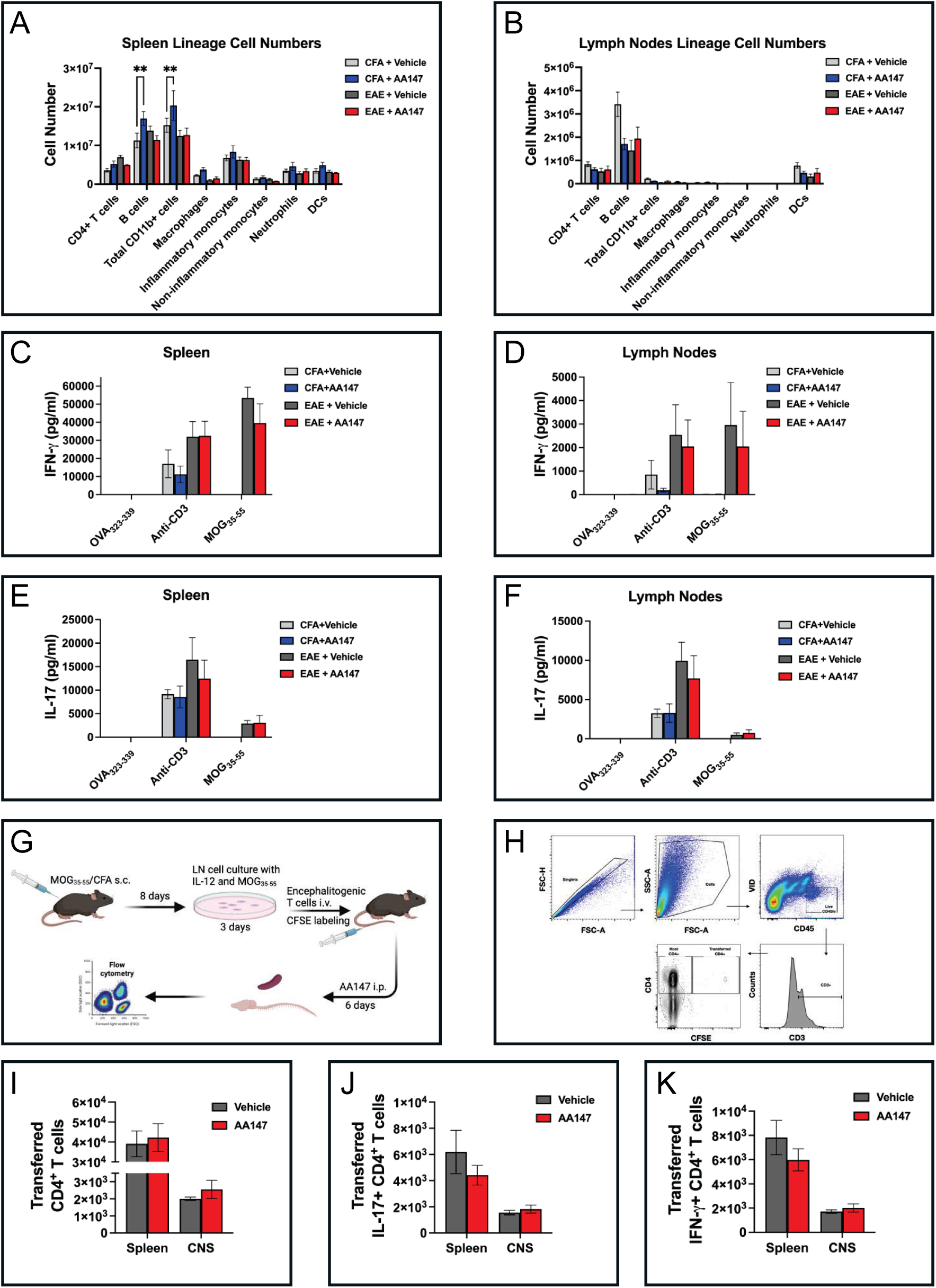
AA147 does not affect the immune cell populations of EAE mice. Splenocytes and lymphocytes were collected from PID12 of CFA or EAE primed mice treated with vehicle or 8mg/kg AA147 beginning from PID7. Flow cytometry analysis for immune cell populations present in the spleen (A) and in the lymph nodes (B) was completed. The level of IFN-γ and IL-17 production was assessed from splenocytes (C,E) and inguinal lymph node cells (D,F) isolated from PID12 of EAE mice. Cells were cultured in the presence of anti-CD3, OVA_323-339_, or MOG_35-55_ for 72 hours. (G-K) The experimental schema for adoptive transfer EAE (G), and gating scheme used for flow cytometric analysis is presented (H). Recipient C57BL/6 mice received 3×10^6^ CFSE-labeled blast cells, and mice received 6 doses either vehicle or AA147. On Day 7 post CFSE-labeled blast cell transfer, CNS and spleen were then collected for flow. The number of transferred (CFSE+) CD4+ T cells (I), IL-17+ transferred CD4+ T cells (J) and IFN-γ+ transferred CD4+ T cells (K) was numerated. Data are expressed as mean ± SEM; n= 5 per group; **p< 0.01 by one-way ANOVA.

To explore the effect of AA147 on T cells trafficking into the CNS, we used the MOG_35-55_ adoptive transfer model of EAE. CFSE-labeled MOG_35–55_ blast cells were transferred into naïve recipient mice that then received daily treatment of either vehicle or AA147 (8mg/kg) (Figure 2G). To determine how AA147 might affect T cell infiltration, spleen and CNS (brain and spinal cord) were collected from recipient mice 7 days post-cell transfer for flow cytometry analysis (Figure 2H). There was no significant difference in the numbers of transferred CD4+ T cells (CFSE labeled), IL-17+CD4+ T cells and IFN-γ+CD4+ T cells in the spleen or CNS between vehicle and AA147 treatment (Figure 2I, J, K).

Collectively, these data indicate that AA147 treatment has no detectable immunosuppressive effect on EAE’s peripheral immune response and trafficking.

### AA147 alters CNS inflammatory response

CNS inflammation in EAE is characterized by infiltration of inflammatory cells, including T lymphocytes and macrophages (Hickey, 1991). We observed a significant reduction in the number of infiltrating immunes cells in the CNS of EAE mice treated with AA147, including CD4+ T cells, B cells, CD11b+ cells and macrophages (p<0.001, p<0.05, p<0.0001, and p<0.05, respectively) (Figure 3A). Analysis of CD4+ T cell subtypes revealed a decrease in T cells expressing Ki67, CD44hi, CTLA4, and IFN (Figure 3B). The significant reduction in B cells was observed solely in Sca-1 positive B cells, a major source of antibody production (Figure 3C). While the overall count of microglia remained constant, there was a notable decrease in MHCII+ microglia in mice treated with AA147, indicating a reduction in activated microglia (p<0.05) (Figure 3D). In line with the flow cytometry study, fewer CD3+ T cells were detected by immunostaining lumbar spinal cord sections from EAE mice treated with AA147 than with vehicle (p<0.05) (Figure 3E, F). In addition, EAE exhibited a significant increase in the number of TMEM119+ activated microglia (p<0.001) as well as CD11b+ macrophages (p<0.01) compared to CFA mice. However, AA147 treatment reduced the activation of microglia (p<0.01) and macrophages (p<0.05) in the lumbar spinal cord (Figure 3G-J). These data demonstrate that AA147 decreases the infiltrating immune cells in the CNS and reduces inflammatory microglia/macrophage activation in EAE mice.

**Figure 3:**
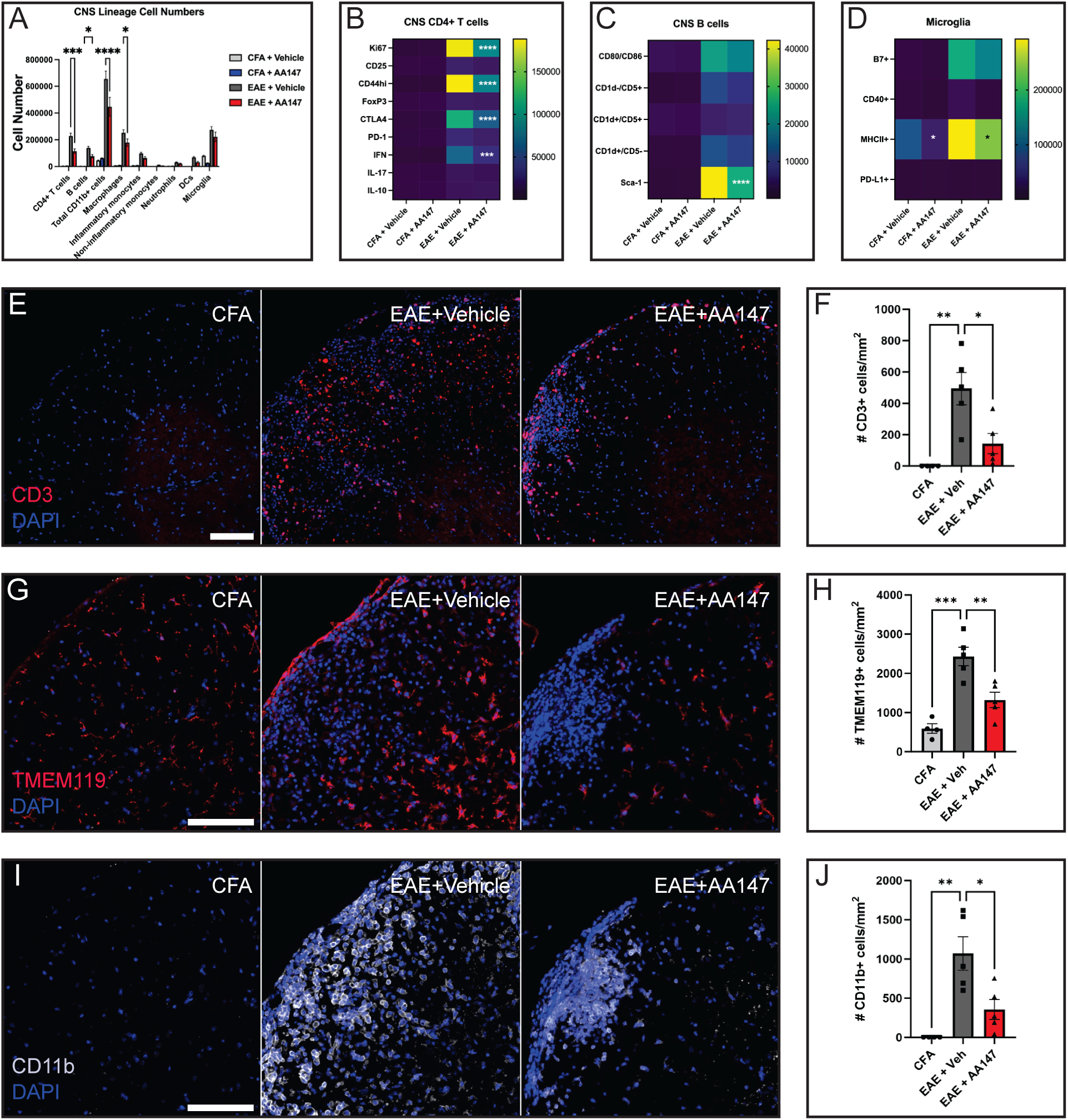
AA147 reduces CNS inflammation of EAE mice. (A-D) Flow cytometry analysis of immune cell populations (A), T cell population (B), B cell population (C) and microglia population (D) in the CNS from PID12 of CFA or EAE mice treated with vehicle or 8mg/kg female AA147 beginning from PID7. Data are expressed as mean ± SEM; n= 5 per group; *p<0.05, ***p< 0.001, ****p< 0.0001 by one-way ANOVA. (E-J) Histology analysis of cross-sections of lumbar spinal cord collected at PID 16 from CFA mice or EAE mice treated daily with vehicle or 8mg/kg AA147. (E) Representative images of CD3 staining (Scale bar=100µm) and (F) quantification of the number of CD3+ T cells in the white matter of lumbar spinal cord. (G) Representative images of TMEM119 staining (Scale bar=100µm) and (H) quantification of the number of TMEM119+ microglia in the white matter of lumbar spinal cord. (I) Representative images of CD11b staining (Scale bar=100µm) and (J) quantification of the number of CD11b+ macrophages in the white matter of lumbar spinal cord. Data are expressed as mean ± SEM; n= 4-5 per group; *p< 0.05, **p< 0.01, ***p<0.001 by one-way ANOVA.

In EAE, the proinflammatory cytokines are critically involved in the initiation and amplification of the local immune response in the CNS, which is counterbalanced by upregulation of anti-inflammatory cytokines (Stoll et al., 2000). We thus examined the impact of AA147 on the cytokine expressions in the CNS of EAE mice using RT-qPCR analysis. Interestingly, we observed that AA147 treatment did not alter mRNA levels of anti-inflammatory cytokines (*Arg-*1, *Il-4* and *Il-10*) in EAE, while it significantly reduced the mRNA levels encoding proinflammatory cytokines (*iNOS*, *Tnf-α* and *Ifn-γ)* (p< 0.05, respectively) (Supplementary Figure 4A-F). There was a trend toward reduced *Il-17* expression, although it was not statistically significant (Supplementary Figure 4G). These data suggest that AA147 treatment results in a relative predominance of an anti-inflammatory response over pro-inflammatory factors.

### AA147 mitigates ER stress and apoptotic cell death

AA147 was initially reported to be a specific ATF6 activator of the UPR (Paxman et al., 2018). Multiple recent studies have suggested that AA147 can also regulate NRF2 in neural-derived cells through an analogous mechanism to that associated with AA147-dependent ATF6 activation (Rosarda et al., 2021; Yuan et al., 2022). Consistent with this, we observed activation of both ATF6 and NRF2 signaling pathways in the CNS of AA147 treated mice (Figure 1A, B). ATF6 signaling plays a key role in regulating ER stress, while NRF2 signaling is crucial for the antioxidative response (Way and Popko, 2016; Motohashi and Yamamoto, 2004). To test whether AA147 could counteract ER-stress-induced apoptosis, we stained lumbar spinal sections with the apoptotic marker anti-cleaved caspase 3. We detected many apoptotic cells (36.73/mm^2^) in the spinal cord of EAE mice treated with vehicle, whereas AA147 significantly reduced apoptotic cell numbers to 9.49 /mm^2^ (p<0.05) (Supplementary Figure 5). Notably, a lower percentage of TPPP+ oligodendrocytes (0.5%) underwent apoptosis, as indicated by cleaved caspase 3 in AA147-treated mice compared to vehicle-treated controls (1.8%, p<0.05), suggesting that the drug may protect these cells from apoptosis (Figure 4A, B). Interestingly, while not statistically significant, we also observed an increased proportion of apoptotic T cells within the CNS of AA147-treated mice compared to vehicle (Figure 4C, D). This trend suggests that the absence of sustained MOG antigen signaling might lead to T cell apoptosis within the CNS in AA147 group. Moreover, we assessed the levels of apoptosis-related proteins as well as the stress markers C/EBP homologous (CHOP) in the lumbar spinal tissues using Western-blots (Figure 4E). As expected, the levels of pro-apoptotic CHOP, BAX, and cleaved caspase 3, significantly increased in EAE compared to CFA (p<0.01, p<0.05, and p<0.05, respectively) (Figure 4F). AA147 treatment inhibited the upregulation of these factors in EAE (p<0.05, p<0.01, p<0.05, respectively) (Figure 4F). To examine whether AA147 could repress oxidative stress, we measured the levels of 3-nitrotyrosine (3-NT), a biomarker of oxidative stress formed due to nitration of tyrosine residues by reactive nitrogen species (Pacher et al., 2007). Western-blots showed that 3-NT levels were significantly higher in EAE+Veh group than in the CFA group (p<0.05). Treatment with AA147 slightly reduced the 3-NT level in EAE, but the reduction was not significant (Figure 4G, H). Similarly, AA147 did not significantly change the level of oxidatively modified proteins in EAE, as detected by dinitrophenylhydrazine (DNPH) derived proteins using the protein carbonyl assay (Figure 4G, I, and Supplementary Figure 6). To further validate these effects, we used a dihydroethidium (DHE) assay to measure the reactive oxygen species (ROS). We did not observe any significant effect of AA147 on ROS generation (Supplementary Figure 7). Collectively, these data suggest that AA147 administered to EAE mice protects oligodendrocytes against cellular stress and apoptosis but has minimal impact on oxidative stress.

**Figure 4:**
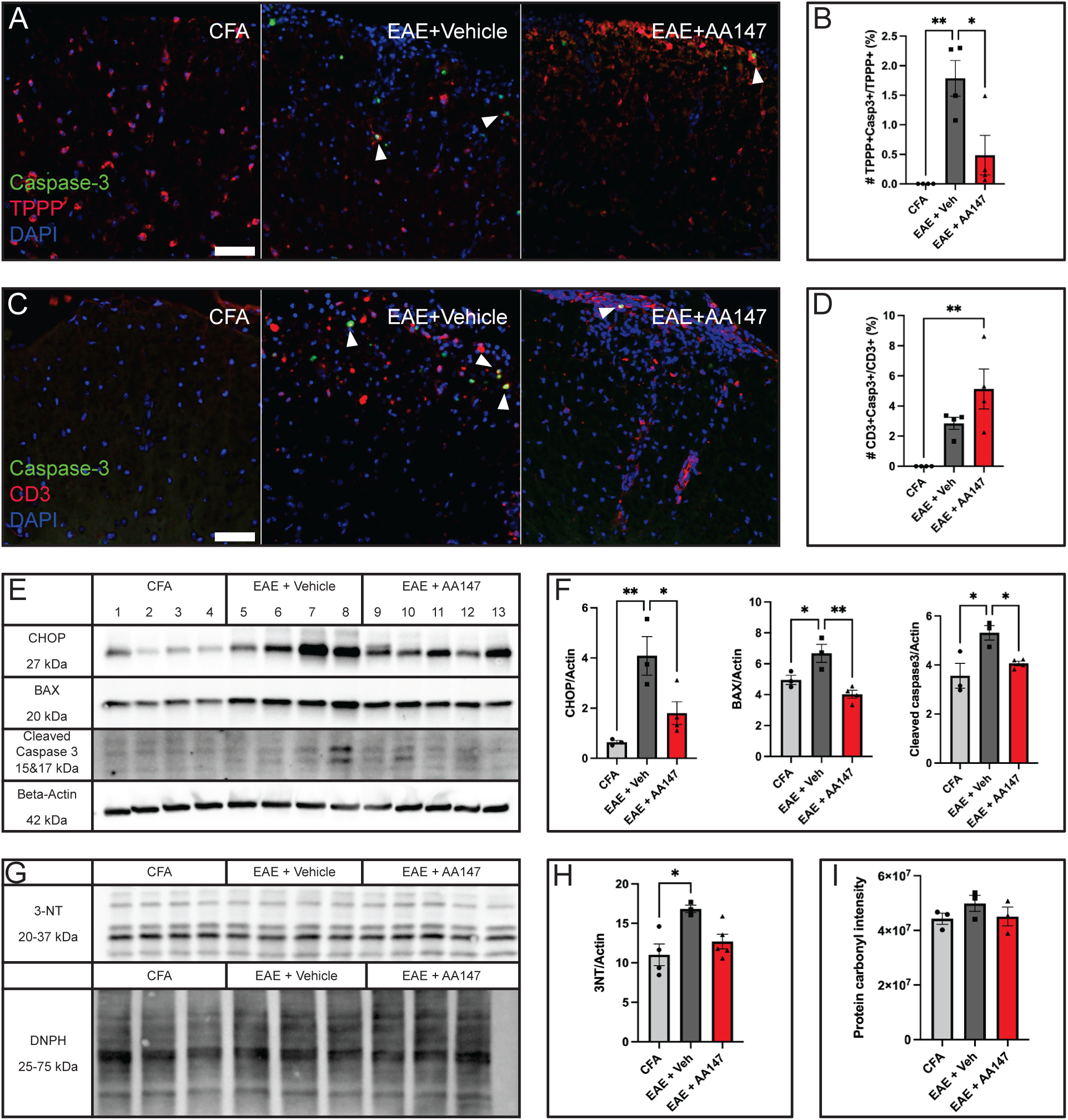
AA147 dampens ER stress induced oligodendrocyte apoptosis but does not significantly affect oxidative stress damage in the CNS of EAE mice. Lumbar spinal cord tissues collected from PID16 of CFA mice or EAE mice treated with vehicle or 8mg/kg AA147 beginning from PID7. (A) Representative images of cleaved caspase 3 and TPPP immunostaining of lumbar spinal cord cross sections (arrow heads point to double positive cells, scale bar=50µm) and (B) the percentage of the number of TPPP+caspase 3+/total TPPP+ cells. (C) Representative images of cleaved caspase 3 and CD3 immunostaining of lumbar spinal cord cross sections (arrow heads point to double positive cells, scale bar=50µm) and (D) the percentage of the number of CD3+caspase 3+/total CD3+ cells. (E) Western blot analysis of stress marker CHOP, apoptosis-related BAX and cleaved caspase 3. (F) Densitometry histograms of CHOP, BAX and cleaved caspase 3 after normalization to β-actin. (G) Western blot analysis of oxidative stress marker, 3NT and the levels of DNPH derived proteins as the markers of protein oxidative damage measured by a protein carbonyl assay kit. (H) Densitometry histograms of 3-NT after normalization to β-actin. (I) Densitometry histograms of protein carbonyl content (DNPH). Data are expressed as mean ± SEM; n= 3-4 per group; *p< 0.05, **p< 0.01 by one-way ANOVA.

### AA147 activates ATF6 and NRF2 signaling pathways in a cell-type-specific manner

AA147 has demonstrated a distinctive potential for activating adaptive ATF6 and/or NRF2 transcriptional signaling in neuronal cells (Rosarda et al., 2021; Yuan et al., 2022). We aimed to determine whether AA147 activates ATF6 and NRF2 pathways in CNS resident cells under inflammatory attack. Activation of these pathways involves translocation to the nucleus to initiate the transcription of target genes. To investigate the subcellular location of endogenous ATF6 and NRF2 in oligodendrocytes, we exposed differentiating oligodendrocytes to IFN-γ, a critical proinflammatory cytokine in MS and EAE, for 16 hours with or without AA147 treatment. A subsequent immunofluorescence study showed that ER membrane bound ATF6 was primarily located in the cytoplasm in the non-stressed control (nontreated, NT) as well as in IFN-γ stressed oligodendrocytes (Figure 5A). Upon AA147 treatment for 16 hours, ATF6 showed increased localization to DAPI-stained nuclei in both NT and IFN-γ stressed cells (Figure 5A). NRF2 was also initially observed in the cytoplasm of NT and IFN-γ stressed cells, but AA147 did not trigger significant nuclear translocation of NRF2 (Figure 5B).

**Figure 5.**
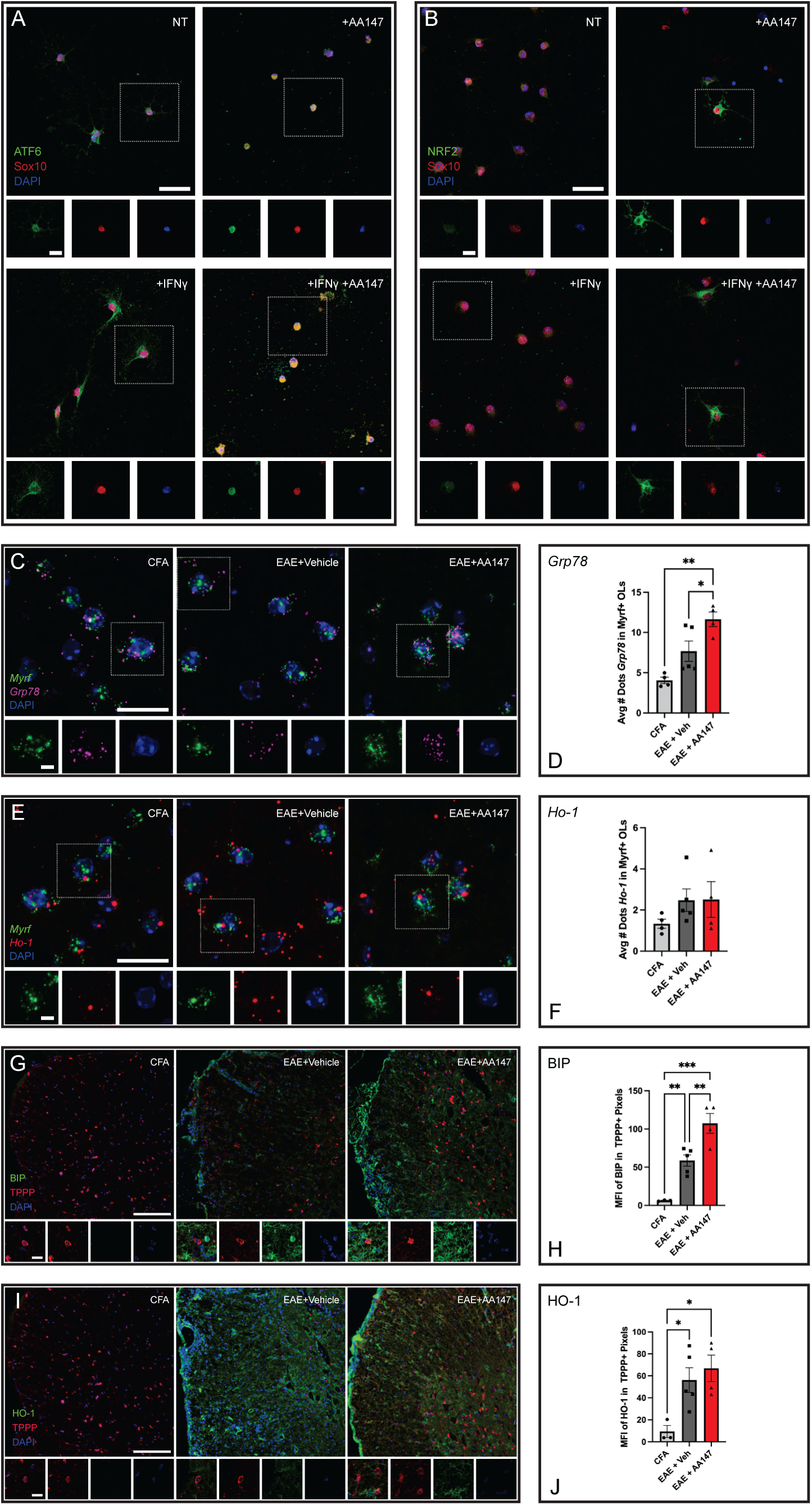
AA147 activates ATF6 pathway, not NRF2 in oligodendrocytes. (A, B) Differentiating oligodendrocytes incubated with AA147 in the presence or absence of IFN-γ. (A) Representative images of cultured oligodendrocytes immunostained with ATF6 and the oligodendrocyte linage marker Sox10 (Scale bar=25µm, 5µm) and (B) stained with NRF2 and Sox10 (Scale bar=25µm, 5µm). (C-J) Lumbar spinal cord tissues collected from PID16 of CFA mice or EAE mice treated with vehicle or 8mg/kg AA147 beginning from PID7. (C) Representative images of ACD RNASCOPE Multiplex v2 staining of lumbar spinal cords using mRNA probes for oligodendrocyte *Myrf* and ATF6 downstream target *Grp78* (Scale bar=20µm, 5µm) and (D) quantification of the number of *Grp78* dots in *Myrf*+ oligodendrocytes. (E) Representative images of ACD RNASCOPE Multiplex v2 staining of lumbar spinal cords using mRNA probes for oligodendrocyte *Myrf* and NRF2 downstream target *Ho-1* (Scale bar=20µm, 5µm) and (F) quantification of the number of *Ho-1* dots in *Myrf*+ oligodendrocytes. (G) Representative images of ATF6 downstream target Grp78/BIP immunostaining (Scale bar=500µm, 10µm) and (H) quantification of average mean fluorescence intensity (MFI) of BIP in TPPP+ oligodendrocytes in the white matter of lumbar spinal cord. (I) Representative images of NRF2 downstream target HO1 immunostaining (Scale bar=500µm, 10µm) and (J) quantification of average MFI of HO1 in TPPP+ oligodendrocytes in the white matter of lumbar spinal cord. Data are expressed as mean ± SEM; n= 4-5 per group; *p< 0.05, **p< 0.01, ***p<0.001 by one-way ANOVA.

Furthermore, we evaluated the expression of *Grp78* mRNA (ATF6 target gene) and *Ho-1* mRNA (NRF2 target gene) in lumbar spinal cord sections from mice subjected to EAE and treated with vehicle or AA147 using RNAscope and immunofluorescent staining. Our RNAscope analysis showed a significant increase in *Grp78* mRNA dots in Myrf+ oligodendrocytes in EAE mice treated with AA147, as compared to vehicle treated EAE mice (p<0.05) (Figure 5C, D). Likewise, the expression of GRP78/BIP protein also increased in TPPP+ mature oligodendrocytes in the EAE+AA147 group, in comparison to the EAE+Veh group (p<0.01) (Figure 5G, H). No difference was detected in *Ho-1* mRNA and protein expression in oligodendrocytes between the vehicle and AA147 groups (Figure 5E, F, I, J). These findings indicate that AA147 activates the ATF6 signaling pathways in inflammatory stressed oligodendrocytes, while the NRF2 signaling pathway remains unaffected.

Microglia are CNS-resident macrophages that are also implicated in the pathogenesis of MS and EAE, as their abundance correlates with the extent of axonal damage in MS lesions (Rasmussen et al., 2007; Miron and Franklin, 2014; Crotti and Ransohoff, 2016). We were thus interested in investigating the effects of AA147 on the microglial response to CNS inflammation. Our immunostaining analysis of lumbar spinal sections from EAE mice revealed a notable increase in GRP78/BIP (p<0.01) and HO-1 expression (p<0.05) in TMEM119+ microglia, as compared to CFA-treated control mice (Figure 6A-D). AA147 further increased GRP78/BIP and HO-1 expression levels in microglia in EAE mice (p<0.05 for both) (Figure 6 A-D). This indicates that AA147 promoted both ATF6 and NRF2 pathways in activated microglia.

**Figure 6:**
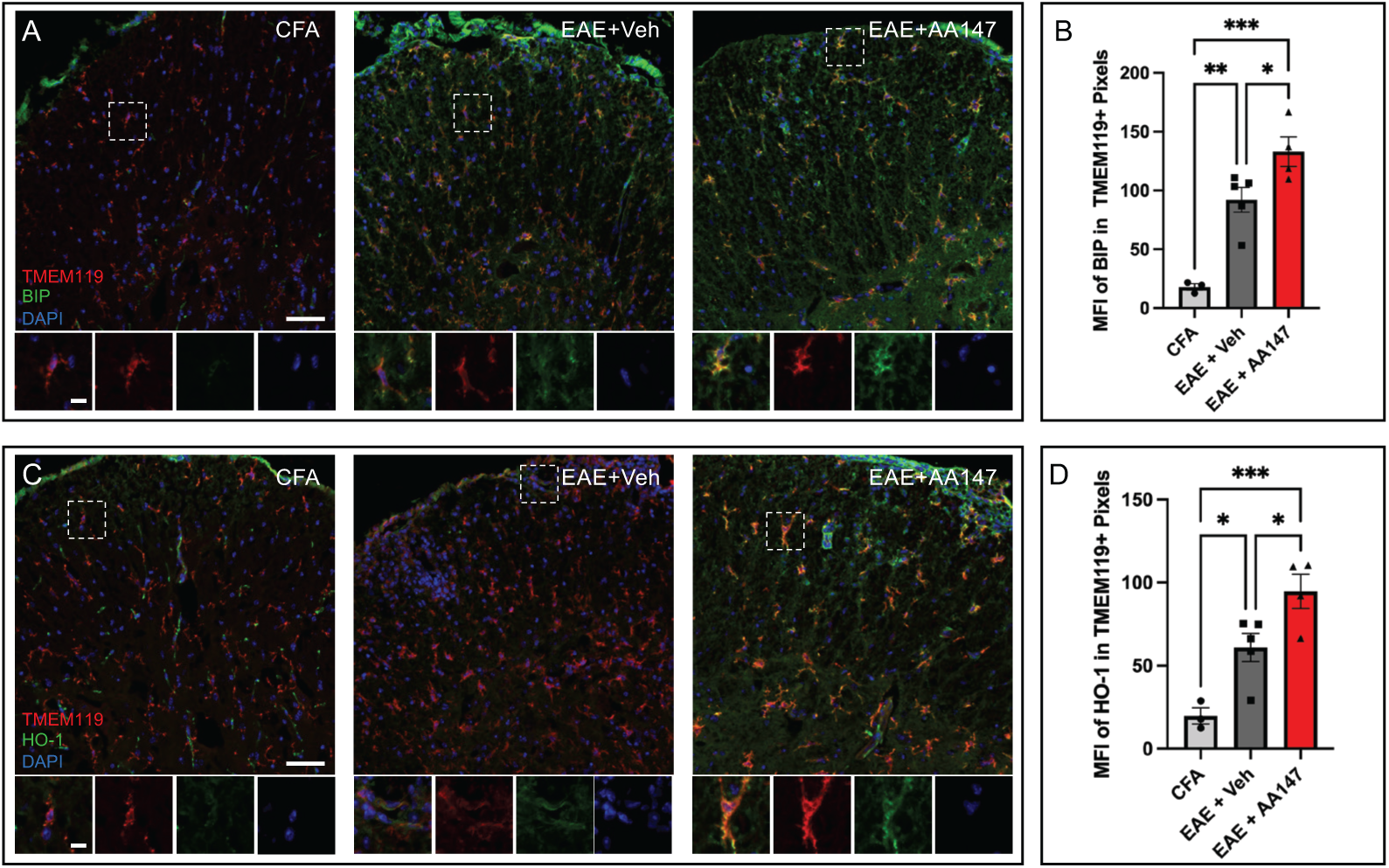
AA147 activates both ATF6 and NRF2 pathways in microglia of EAE mice. Histology analysis of cross sections of lumbar spinal cord collected at PID 16 from CFA mice or EAE mice treated daily with vehicle or 8mg/kg AA147. (A) Representative images of GRP78/BIP and microglia marker TMEM119 immunostaining (Scale bar=50µm, 10µm) and (B) quantification of MFI of BIP in TMEM119+ microglia. (C) Representative images of HO-1 and TMEM119 immunostaining (Scale bar=50µm, 10µm) and (D) quantification of MFI of HO-1 in TMEM119+ microglia in the white matter of lumbar spinal cord. Data are expressed as mean ± SEM; n= 3-5 per group; *p< 0.05, **p< 0.01, ***p<0.001 by one-way ANOVA.

### AA147 changes microglia profiling during the disease course

Our results suggest that AA147 activates the ATF6 and NRF2 pathways in microglia but does not affect oxidative stress levels. Recent studies have demonstrated that pharmacological activation of NRF2 by sulforaphane can induce the Arginase1 (ARG1) + microglia phenotype in vitro and in chronically stressed animals (He et al., 2022; Tang et al., 2022). Microglia activation is heterogeneous, and the ARG1+ microglia cluster is essential for proper brain development (Stratoulias et al., 2023). ARG1 is an enzyme converting arginine into ornithine and urea. iNOS competes for the same substrate, arginine, to produce NO (Quirino et al., 2013). NO promotes the synthesis of IL-6 and other inflammatory factors, all of which are involved in the microglia inflammatory response (Bogdan, 2015). Thus, to investigate whether AA147 alters microglia phenotypes, we conducted immunostaining on the lumbar spinal cord sections using anti-ARG1 and anti-iNOS antibodies. Our data shows that EAE mice significantly increased ARG1 immunoreactivity in TMEM119+ microglia when compared to CFA (p<0.01). AA147 treatment further enhanced ARG1 expression in microglia of EAE mice (p<0.05) (Figure 7A, B). Likewise, EAE mice presented a significantly higher iNOS immunoreactivity in microglia than CFA (p<0.001), but AA147 did not alter iNOS expression in EAE microglia (Figure 7C, D). Collectively, these data demonstrate that AA147 promotes ARG1+ phenotypes of microglia possibly through activating the ATF6 or NRF2 pathways during EAE progression.

**Figure 7:**
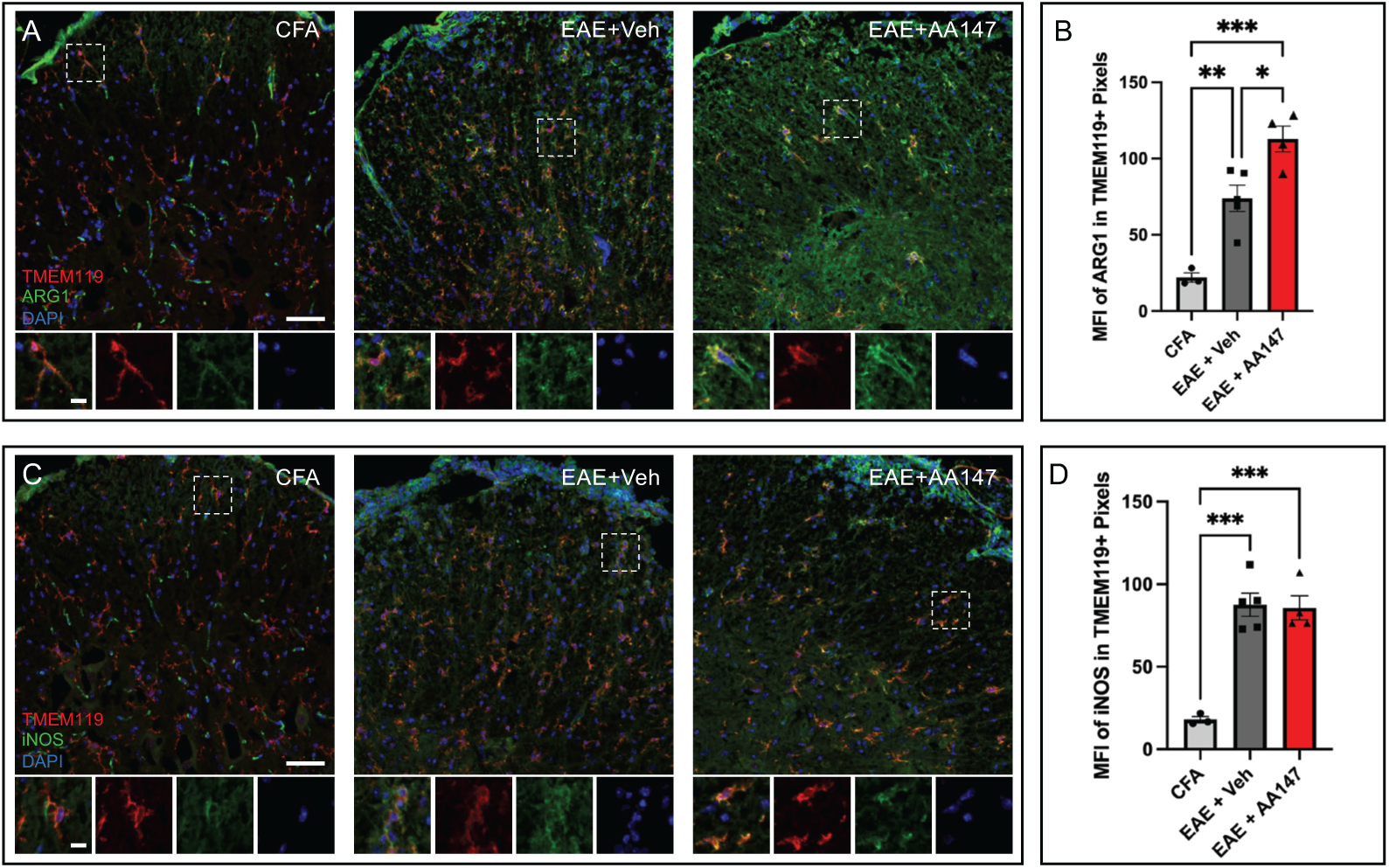
AA147 increases ARG1 levels in microglia of EAE mice. Histology analysis of cross-sections of lumbar spinal cord collected at PID 16 from CFA mice or EAE mice treated daily with vehicle or 8mg/kg AA147. (A) Representative images of ARG1 and microglia marker TMEM119 immunostaining (Scale bar=50µm, 10µm) and (B) quantification of MFI of ARG1 in TMEM119+ microglia. (C) Representative images of iNOS and TMEM119 immunostaining (Scale bar=50µm, 10µm) and (D) quantification of MFI of iNOS in TMEM119+ microglia in the white matter of lumbar spinal cords. Data are expressed as mean ± SEM; n= 3-5 per group; *p< 0.05, **p< 0.01, ***p<0.001 by one-way ANOVA.

### Deficiency of ATF6 in oligodendrocytes abrogates the protective effect of AA147 on EAE

To clarify the cellular specificity of the ATF6 signaling pathway on oligodendrocytes in EAE mice, we employed a conditional and tamoxifen-inducible *Atf6alpha* knockout mouse line (*Atf6^fl/fl^; PLP-CreER^t^*) to inactivate ATF6 in adult oligodendrocytes. Both *Atf6^fl/fl^; PLP-CreER^t^*+ and *Atf6^fl/fl^; PLP-CreER^t^*-control mice were given IP injections of tamoxifen three weeks before EAE induction. In the presence of PLP-Cre (*Atf6^fl/fl^;PLP-CreER^t^*+), ATF6 expression is remarkably diminished in TPPP+ oligodendrocytes (Figure 8A, B; Supplementary Figure 8), while remaining consistently observable in TMEM119+ microglia and GFAP+ astrocytes, as evidenced by immunostaining of spinal cord sections (Figure 8C, D). Next, we found that *Atf6^fl/fl^;PLP-CreER^t^*+ mice displayed a similar EAE disease progression of *Atf6^fl/fl^;PLP-CreER^t^*-, suggesting that knocking out ATF6 only in oligodendrocytes does not affect EAE severity (Figure 8E). To determine if the protective effect of AA147 is mediated by enhancing ATF6 signaling in oligodendrocytes, we treated *Atf6^fl/fl^;PLP-CreER^t^*+ and *ATF6fl/fl; PLP-CreER^t^-* mice with AA147 starting from PID 7. As expected, AA147 significantly reduced disease severity in the control mice (*Atf6^fl/fl^;PLP-CreER^t^-*) (p<0.05 and p<0.01) (Figure 8E). Nevertheless, this protection of AA147 was abrogated when ATF6 was deleted in oligodendrocytes from *Atf6^fl/fl^;PLP-CreER^t^*+ mice at the early stage of EAE. In the chronic stage of EAE beyond 30 days, AA147 commenced to alleviate EAE disease in *Atf6^fl/fl^;PLP-CreER^t^*+ mice (p<0.05). These findings indicate that the role of ATF6 in oligodendrocytes is crucial in mediating the protective effects of AA147 against EAE, particularly during the initial stages of disease progression.

**Figure 8:**
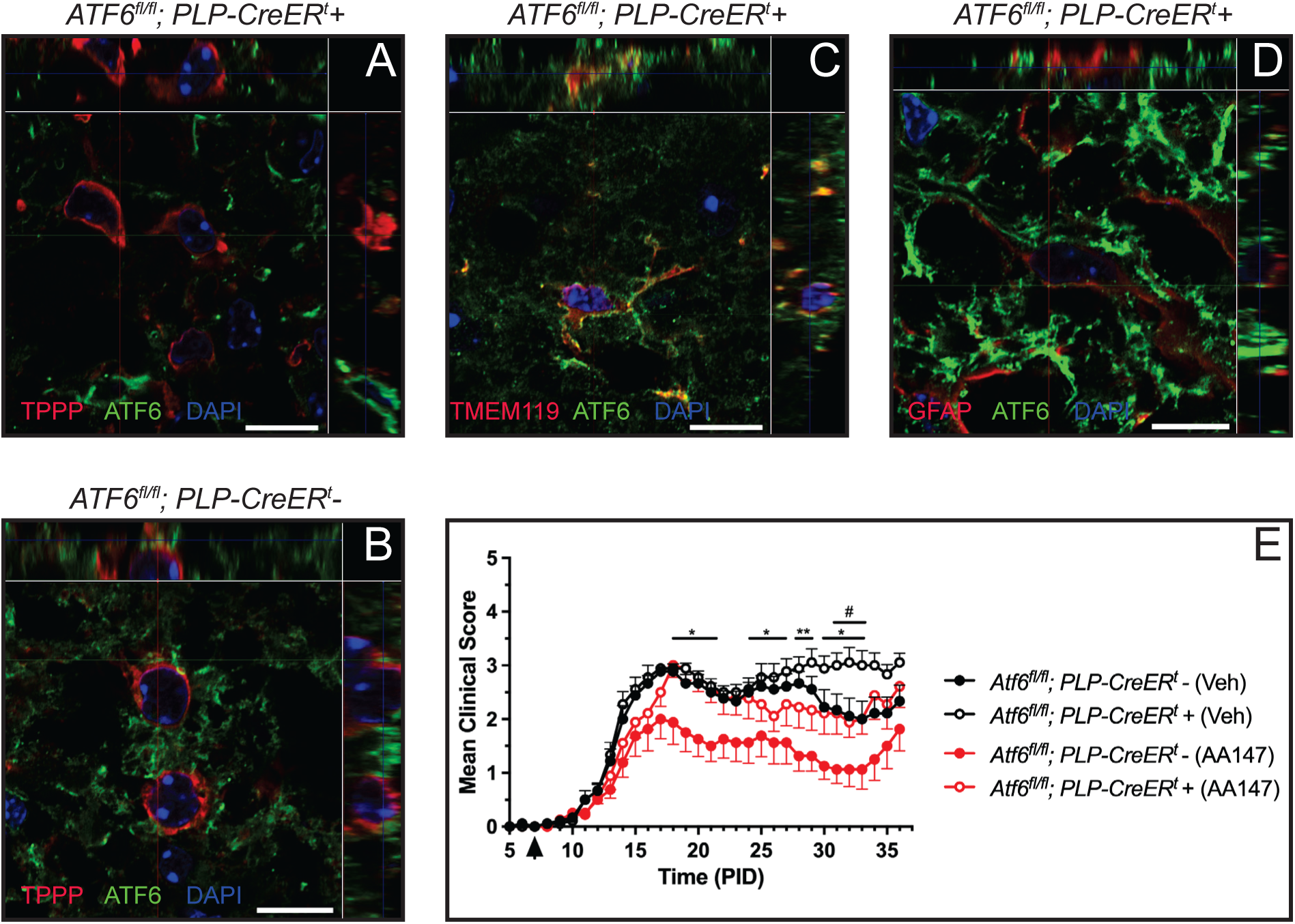
AA147’s protective effect is abrogated in the acute stage of EAE mice lacking ATF6 in oligodendrocytes. (A-D) Histology analysis of cross sections of lumbar spinal cord collected from *ATF6 ^fl/fl^;PLP-CreER^t^-* and *ATF6 ^fl/fl^;PLP-CreER^t^+ mice.* Top and right in each panel show orthogonal view of different planes (X, Y, Z) at a confocal plane. Representative images of oligodendrocyte marker TPPP and ATF6 immunostaining in lumbar spinal cords of *ATF6 ^fl/fl^;PLP-CreER^t^-* mice (A) and *ATF6 ^fl/fl^;PLP-CreER^t^+ mice* (B). Representative images of microglia marker TMEM119 and ATF6 immunostaining (C) and astrocytes marker GFAP and ATF6 immunostaining (D) of lumbar spinal cords of *ATF6 ^fl/fl^;PLP-CreER^t^+ mice.* All scale bars =10µm. (E) Clinical scores of *ATF6 ^fl/fl^;PLP-CreER^t^-* and *ATF6 ^fl/fl^;PLP-CreER^t^*+ female mice immunized with MOG_35-55_/CFA to induce EAE, treated with vehicle and 8mg/kg AA147, starting from PID 7 (arrow pointed). n= 9 per group; *p< 0.05, **p< 0.01 (Cre-with Veh vs Cre-with AA147), ^#^p<0.05 (Cre+ with Veh vs Cre+ with AA147) by two-way ANOVA.

## DISCUSSION

The protective effect of overexpressing of the active ATF6 domain observed in cellular and mouse models of various diseases, including protein-misfolding diseases and stroke, highlights the unique potential for targeting ATF6 to intervene in human diseases (Kaneko et al., 2010; Yu et al., 2017). In the present study, we showed that AA147, an ATF6 activator compound, dampened the clinical symptoms in EAE, protected oligodendrocyte and myelin, reduced axon degeneration, and mitigated CNS inflammation without affecting the peripheral immune response. We also found that AA147 exhibited CNS protective effects partially by suppressing ER stress and apoptosis. Furthermore, in vitro and in vivo studies revealed differential effects of AA147 on the ATF6 and NRF2 pathways in oligodendrocytes and microglia. Lastly, we demonstrated that the protective effect of AA147 was abrogated in the early stage of EAE by inactivating *Atf6* in oligodendrocytes. Together, these data demonstrate that AA147 protects against EAE by shielding oligodendrocytes from inflammatory stress and promoting microglia phenotype shifting, highlighting the potential for AA147 as an MS treatment.

The UPR is an adaptive program that aims to either restore protein homeostasis or trigger cell death upon irreversible stress (Clayton and Popko, 2016). Accumulating evidence suggests that targeting the UPR could offer promising therapeutic avenues for managing MS (Way and Popko, 2016; Lin and Stone, 2020). Small molecules that selectively prolong the PERK branch activation of the UPR, such as Guanabenz and its derivative Sephin1, have been found to delay EAE disease onset and reduce oligodendrocyte death (Way et al., 2015; Chen et al., 2019, 2023). However, research into the potential impact of ATF6 activation on MS remains limited. AA147, the newly identified ATF6 activator, has been shown to provide protection against ischemia/reperfusion damage to various tissues and induce neuroprotection following post-cardiac arrest (Blackwood et al., 2019; Shen et al., 2021; Yuan et al., 2022). Here, we examined whether AA147 confers CNS protection in the MS animal model, EAE.

According to a previous time-course experiment, ATF6-regulated gene induction peaked at 24 hours after AA147 administration (Blackwood et al., 2019). We therefore administered daily IP injections to maintain sufficient drug levels during the disease course. No side effects were detected in mice treated for extended periods using this regimen. We noted that AA147 treatment initiated immediately before disease onset significantly alleviated EAE disease progression, reduced CNS inflammation, increased oligodendrocyte survival and preserved myelin/axon integrity. Reduced apoptotic cell death of oligodendrocytes was observed in AA147 treated mice. However, it is worth noting that less than 10% of caspase 3+ cells were oligodendrocytes in EAE+Veh, suggesting that AA147 rescues oligodendrocytes partially through reduced apoptosis. Given that previous studies indicate that oligodendrocyte death in MS and EAE is likely driven by other mechanisms (Bonetti and Raine, 1997; Prineas et al., 2001), additional studies investigating non-apoptotic pathways could provide more insights how AA147 supports oligodendrocyte resilience.

During EAE, autoreactive T cells infiltrate the CNS and are reactivated by APCs displaying CNS self-antigen (Goverman, 2009). Our data indicate that AA147 neither affects the activation of T cells in the periphery nor T cell trafficking into the CNS, suggesting that AA147 could confer protection to the CNS by limiting myelin debris as epitope antigens and thereby reducing T cell reactivation in the CNS (Titus et al., 2020). A similar CNS target effect, rather than peripheral target, was also reported for another UPR activator Sephin1 in our previous study (Chen et al., 2019). The success of this preventive treatment encourages further investigation into the effects of therapeutic regimens, particularly treatment initiation at the disease onset or even peak, which would further underscore AA147’s clinical significance.

AA147 was among the first small molecules identified as specific ATF6 activators through phenotypic screening (Plate et al., 2016; Paxman et al., 2018). Blackwood et al. showed that AA147 treatment induces the expression of the ATF6 target genes in the heart, but not PERK-or IRE1-regulated genes (Blackwood et al., 2019). In agreement, our study also revealed an ATF6 pathway-specific activation by AA147 in the CNS. It is worth noting that there is coordination and interdependency between the ATF6 and IRE1 pathways. Under ER stress, ATF6 promotes the transcription of the *Xbp1* gene, which is then spliced by IRE1 to produce active XBP1 mRNA (Yoshida et al., 2001; D’Amico et al., 2022). Our study found no evidence of Xbp1 splicing when treated with AA147 (Supplementary Figure 1C), further indicating that enhanced ATF6 activation by AA147 does not influence IRE1/XBP1 pathway in the CNS.

A previous study showed that ATF6 deficiency increased disease severity, and exacerbated oligodendrocyte death and demyelination in EAE mice (Stone et al., 2018). Conversely, another study suggested that deletion of ATF6 ameliorates demyelination and disease severity in EAE mice by inhibiting microglia-mediated inflammation (Ta et al., 2016). Although the different EAE immunization regimens could account for the disparities between these studies, the exact role of ATF6 in EAE mice, especially in specific cell types, remains unclear. In the present study, we induced EAE in oligodendrocyte-specific-*Atf6* mutant mice. Surprisingly, no difference on clinical presentation was detected when ATF6 was missing from oligodendrocytes, indicating that ATF6 pathway is not activated in oligodendrocytes facing inflammatory attack. This data also suggests that increased EAE severity in ATF6 null mice (Stone et al., 2018) was likely due to cells other than oligodendrocytes.

Although the absence of ATF6 did not make the oligodendrocytes vulnerable, the beneficial effect of AA147 before PID30 was abolished in EAE mice with an ATF6 defect in oligodendrocytes. This suggests that AA147 affords protection during the acute phase of disease by enhancing the ATF6 pathway in oligodendrocytes, thereby increasing their resilience to stress. These findings are consistent with previous studies showing that enhanced ATF6 activity can be protective in various disease models (Grandjean and Wiseman, 2020). However, in the chronic phase of EAE (after PID30), the therapeutic effect of AA147 was observed in mice lacking ATF6 in oligodendrocytes, suggesting that AA147 might also activate additional protective mechanisms in other cell types to mitigate disease pathology.

To examine this issue, we explored additional cytoprotective pathways and resident cells in the CNS. It has been reported that AA147 activates NRF2 selectively in neuronal-derived cell lines (Rosarda et al., 2021). In our study, we found that AA147 activated both the ATF6 and NRF2 pathways in microglia. NRF2 is known to induce an antioxidative stress response upon activation by inducing antioxidant and detoxifying enzymes (Motohashi and Yamamoto, 2004). In contrast to a previous study where AA147 reduced oxidative stress induced toxicity in several cell types by decreasing ROS-associated damage (Blackwood et al., 2019), we were surprised to find that AA147 did not reduce oxidative stress in the CNS of EAE mice. There are two possible explanations: 1) the level of increased NRF2 by AA147 is not enough to generate a sufficient antioxidative response to defend against oxidative stress; 2) other NRF2-regulated responses, particularly in microglia, could be at play.

Recent evidence from studies on depression-like mice indicate that NRF2 activation can program microglia to an ARG1+ phenotype (He et al., 2022; Zhang et al., 2023). Consistent with this, our findings demonstrate that AA147 promotes the expression of ARG1 more prominently than iNOS, possible by regulating the NRF2/HO-1 pathway in microglia (Kan et al., 2015). Traditionally, microglial iNOS is associated with the immune response’s proinflammatory phase, while ARG1 has been linked with the anti-inflammatory phase. However, it is increasingly recognized that iNOS and arginase are often co-expressed, and evidence from single-cell RNA-seq studies suggests that microglia “activation” is a dynamic response involving transcriptionally and spatially distinct subpopulations, challenging the concept of microglial polarization (Hammond et al.; Kim et al., 2016; Ransohoff, 2016). A recent postnatal development study found that ARG1+ microglia have more phagocytic inclusions than ARG1-microglia. Some inclusions contain structures with synaptic vesicles, indicating a phagocytic signature and possible synaptic modification during early postnatal mouse development (Stratoulias et al., 2023). Given the diversity of activated microglia subpopulations found in mouse demyelinated lesions and human MS (Hammond et al., 2019), further research is needed to investigate whether and how the arginase expressed in microglia regulates neuroprotection during CNS inflammation.

In summary, our study demonstrates that AA147 significantly dampens disease severity in mouse models of MS. It is effective in protecting oligodendrocytes from ER stress, preserving myelin from inflammatory attacks, and modulating microglia profiling. These beneficial effects suggest that AA147 is a promising therapeutic for inflammatory demyelinating disorders such as MS.

## DATA AVAILABILITY

The data that support the findings of this study are available from the corresponding author, upon reasonable request.

## Acknowledgement

We thank Joseph Schluep, Erica Garcia, Samantha Wills and Peyton Fay from Loyola University and Young Hyun Che from Northwestern University for technical assistance. This study was supported by a National Multiple Sclerosis Society Career Transition Fellowship TA-2008-37043 (YC), the Dr. Miriam and Sheldon G. Adelson Medical Research Foundation (BP), NIH NINDS 1R35NS137478 (BP) and the National Institutes of Health (AG046495; RLW, JWK).

