## Supplemental Information for "AA147 Alleviates Symptoms in a Mouse Model of Multiple Sclerosis by Reducing Oligodendrocyte Loss"

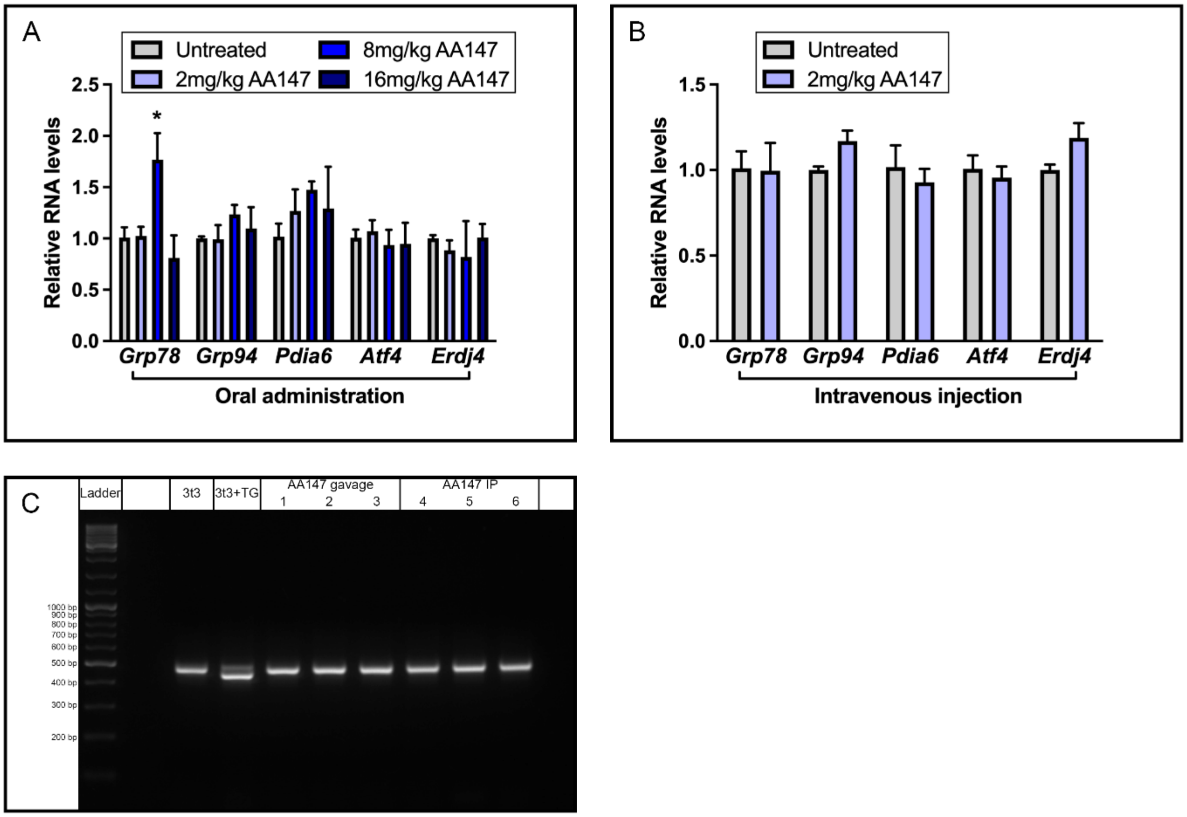


**Supplementary Figure 1:** Relative mRNA levels of UPR genes in lumbar spinal cord of C57BL/6J mice 24h after AA147 treatment. Total RNA was isolated from lumbar spinal cord tissue of C57BL/6J female mice 24 hours after administrating AA147. Relative mRNA expressions of UPR genes were determined by qRT-PCR after oral administration (gavage) (A) or intravenous injection (B). Data are expressed as mean ± SEM; n=3 per group; *$p < 0.05$ by one-way ANOVA. *Xbp1* mRNA splicing was determined by semi-quantitative RT-PCR splicing assay after oral administration (gavage) and IP injection of 8mg/kg AA147. 3T3 cells line in the present or absence of thapsigargin (TG) worked as negative and positive control, respectively (C).

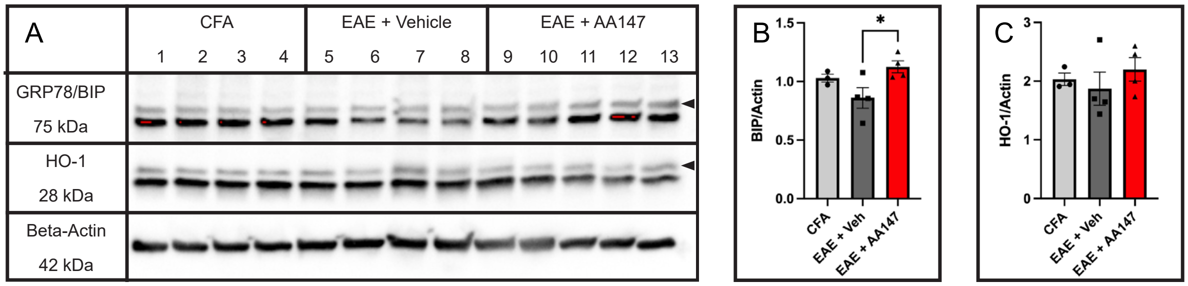


**Supplementary Figure 2: BIP and HO-1 protein levels in the lumbar spinal cord of EAE mice treated with AA147.** Lumbar spinal cord tissues collected from PID16 of CFA mice or EAE mice treated with vehicle or 8mg/kg AA147 beginning from PID7. (A) Western blot analysis of GRP78/BIP (arrowhead pointed) and HO-1(arrowhead pointed) and (B) densitometry histograms of GRP78/BIP and (C) HO-1 after normalization to β-actin. Data are expressed as mean ± SEM; n=3-4 per group; *$p < 0.05$ by one-way ANOVA.

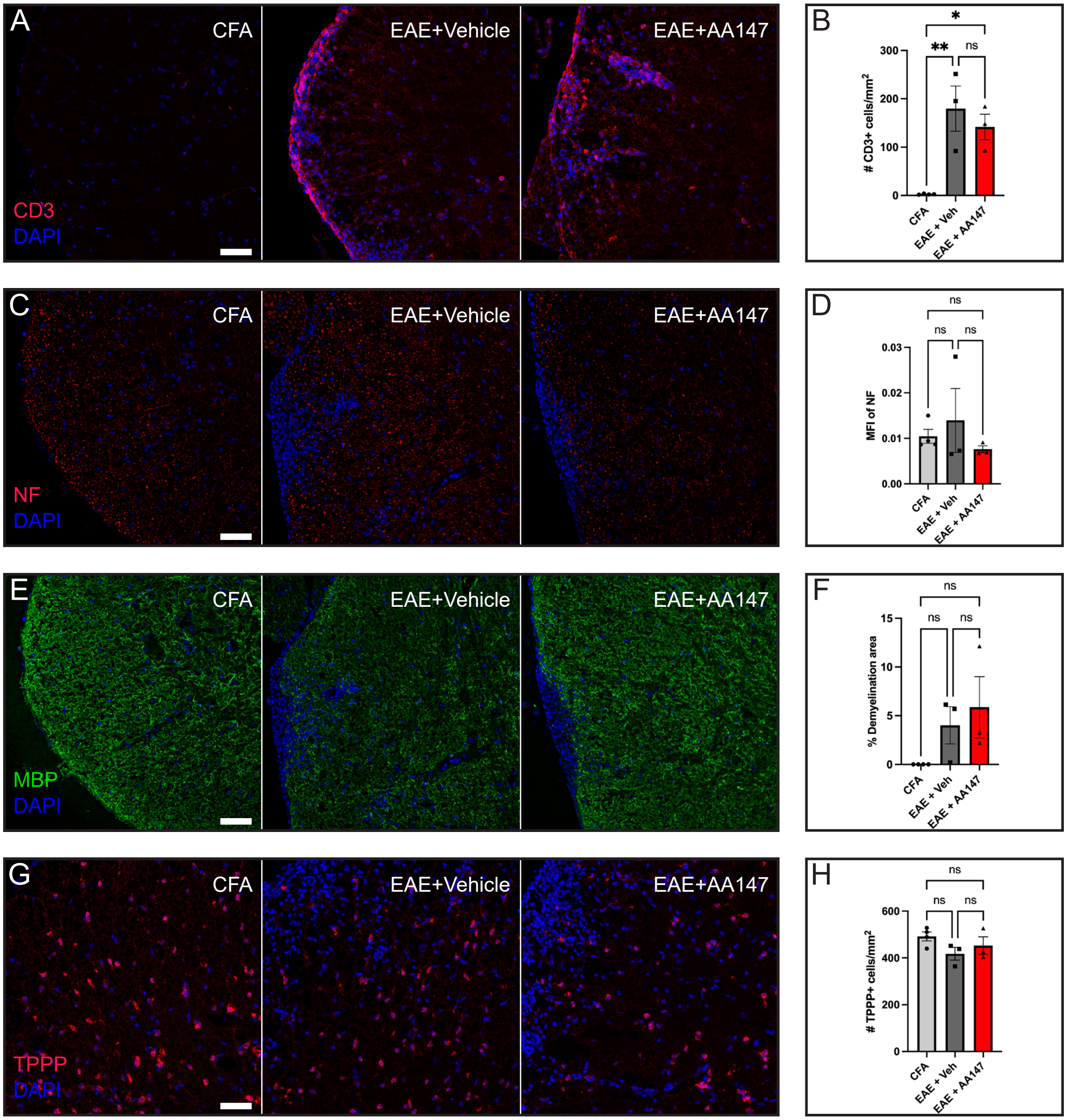


**Supplementary Figure 3: AA147 does not affect CNS inflammation and neuropathology in the early stage of EAE.** Histology analysis of cross sections of lumbar spinal cord collected at PID 12 from CFA mice or EAE mice treated daily with IP injection of vehicle or 8mg/kg AA147. (A) Representative images of CD3 staining (Scale bar=50µm) and (B) quantification of density of CD3 in the white matter of lumbar spinal cord. (C) Representative images of NF staining (Scale bar=50µm) and (D) MFI of NF in the white matter. (E) Representative images of MBP staining (Scale bar=50µm) and (F) the percentage of demyelinated area in the white matter. (G) Representative images of TPPP staining (Scale bar=50µm) and (H) the number of TPPP+ oligodendrocytes in the white matter of lumbar spinal cord. Data are expressed as mean ± SEM; n= 3 per group; *p< 0.05, **p< 0.01 by one-way ANOVA.

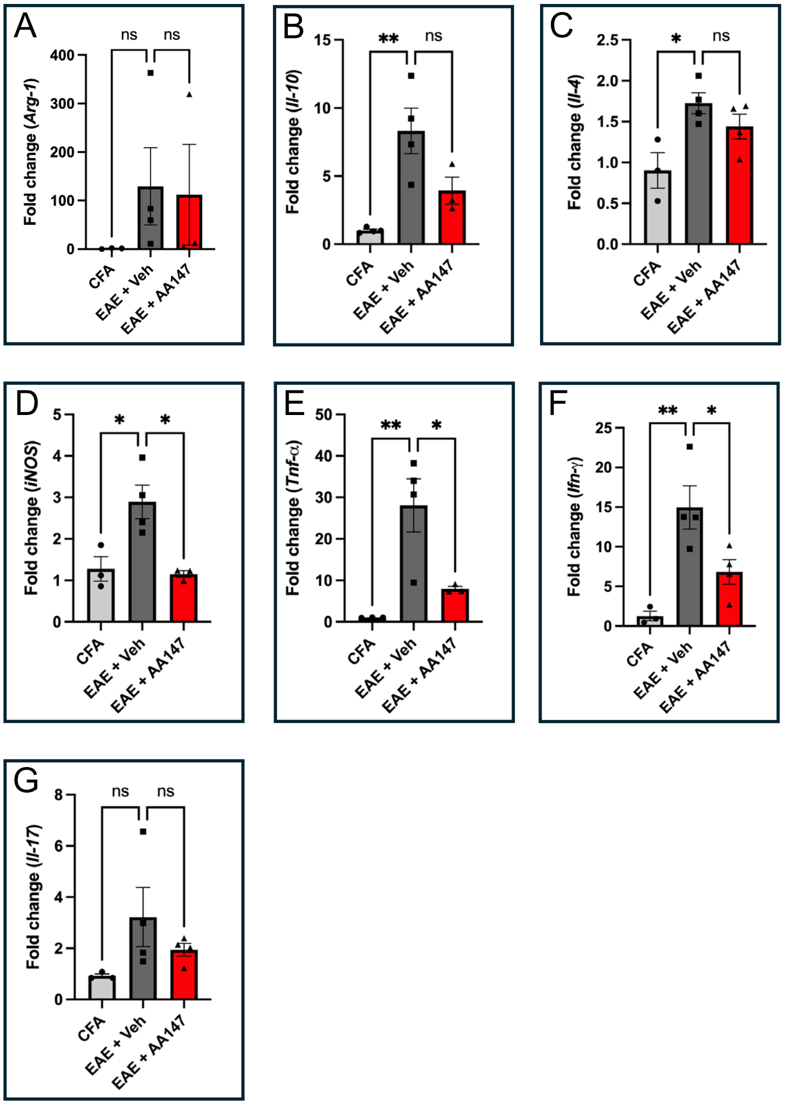


**Supplementary Figure 4: Relative mRNA levels of cytokines in the lumbar spinal cord of EAE mice treated with AA147.** Total RNA was isolated from lumbar spinal cord tissue of PID16 of CFA mice or EAE mice treated with vehicle or 8mg/kg AA147 beginning from PID7. Relative mRNA expressions of *Arg1*(A), *Il-10* (B), *Il-4* (C), *iNOS* (D), *Tnf-α* (E), *Ifn-γ* (F) and *Il-17* (G) were determined by qRT-PCR. Data are expressed as mean ± SEM; n=3-4 per group; *p< 0.05, **p<0.01 by one-way ANOVA.

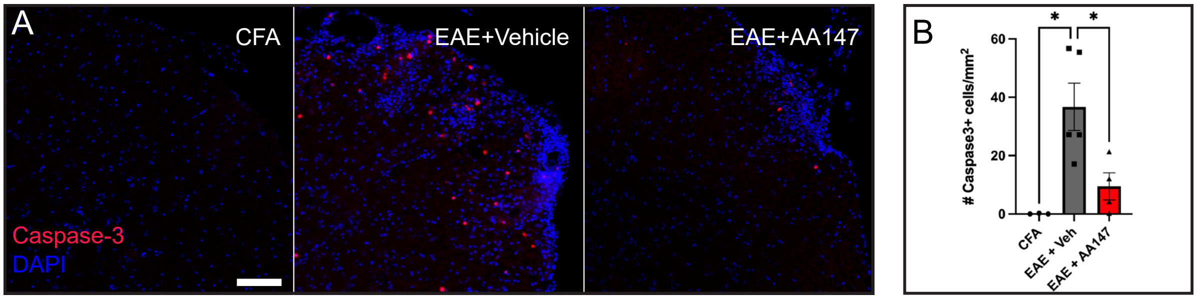


**Supplementary Figure 5: AA147 reduced apoptotic cell death in the lumbar spinal cord of EAE mice.** Lumbar spinal cord tissues collected from PID16 of CFA mice or EAE mice treated with vehicle or 8mg/kg AA147 beginning from PID7. (A) Representative images of cleaved caspase 3 immunostaining of lumbar spinal cord cross sections (Scale bar=100µm) and (B) quantification of the number of caspase 3+ apoptotic cells.

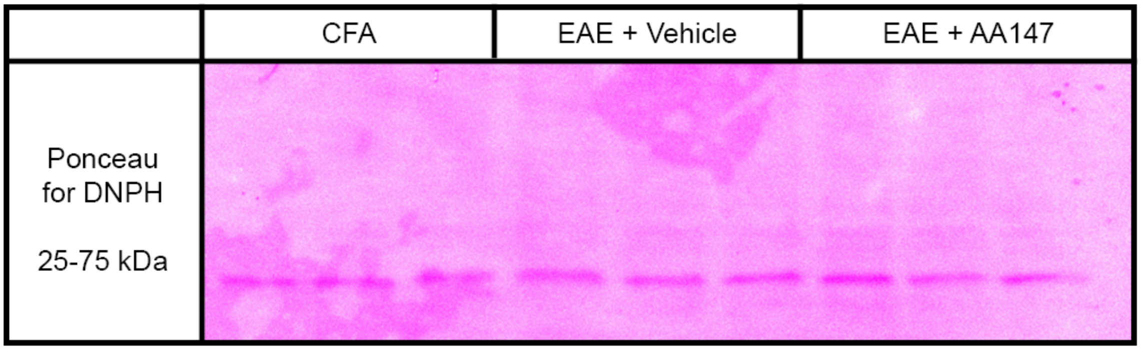


**Supplementary Figure 6:** Ponceau staining of DNPH blot. Lumbar spinal cord tissues collected from PID16 of CFA mice or EAE mice treated with vehicle or 8mg/kg AA147 beginning from PID7. DNPH derived proteins were run on SDS-PAGE and transferred to nitrocellulose membrane with Ponceau staining.

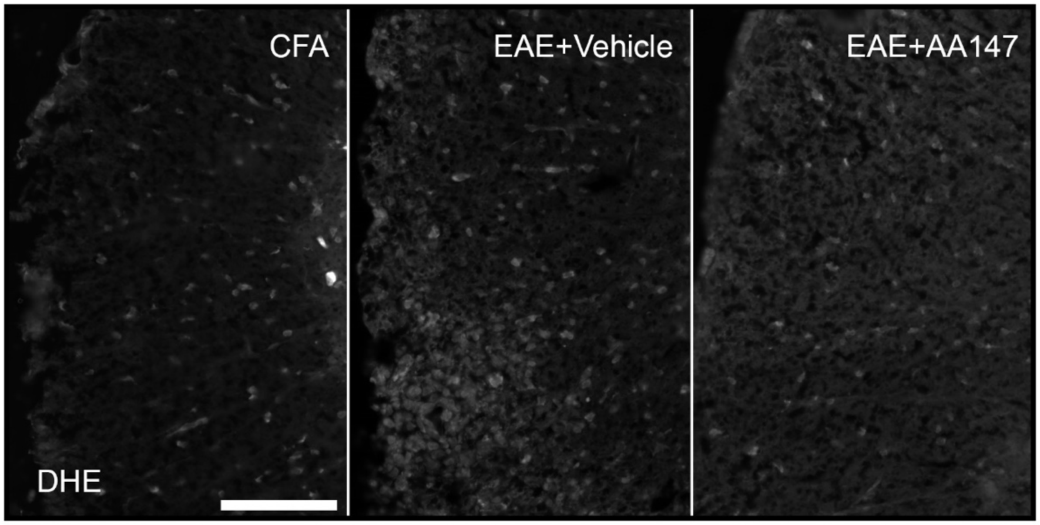


**Supplementary Figure 7:** DHE staining of lumbar spinal cord sections. Lumbar spinal cord tissues collected from PID16 of CFA mice or EAE mice treated with vehicle or 8mg/kg AA147 beginning from PID7. Scale bar=100μm.

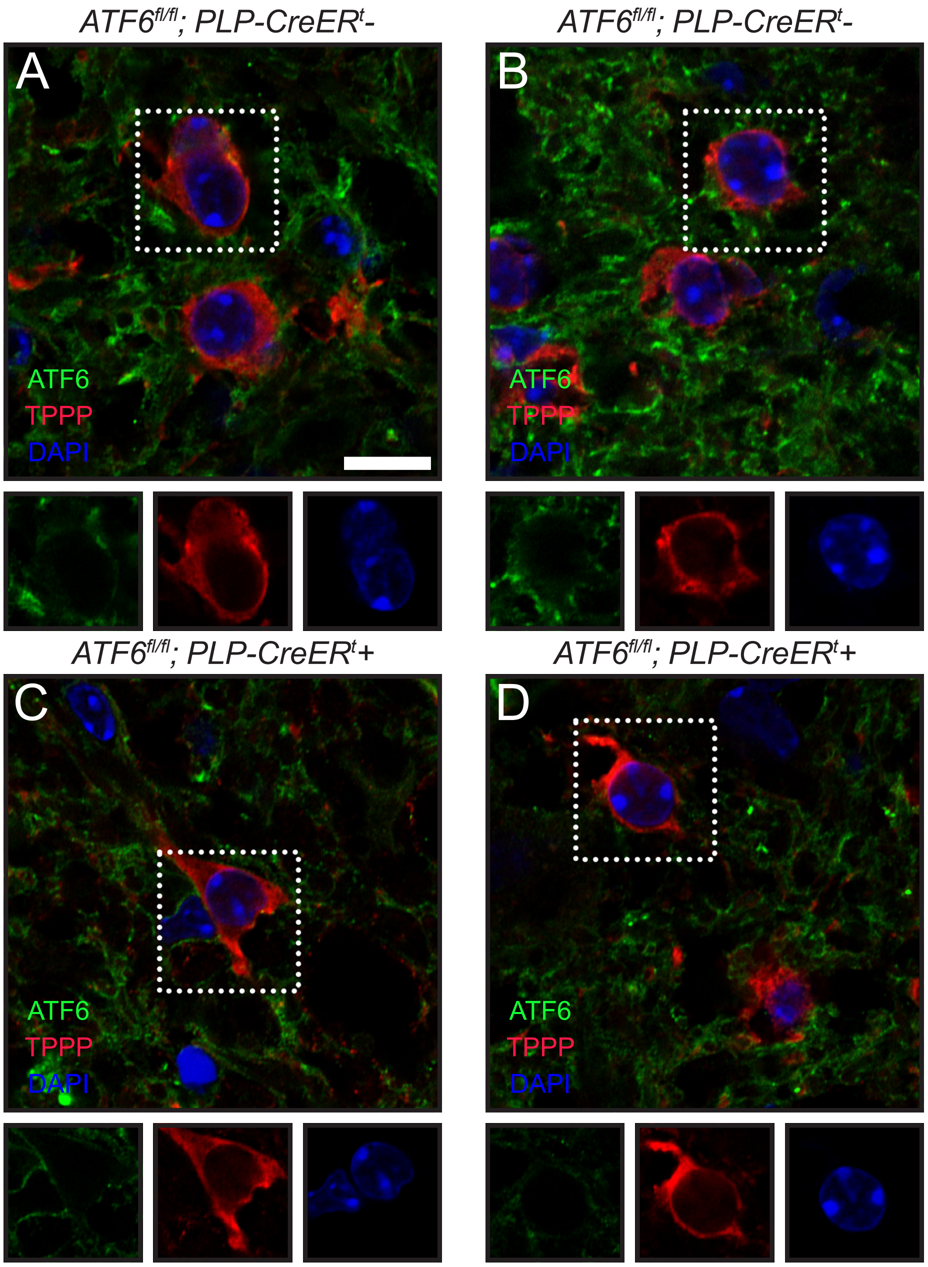


**Supplementary Figure 8: ATF6 expression is diminished in oligodendrocytes in *ATF6 fl/fl; PLP-Cre+ mice.*** Histology analysis of cross sections of lumbar spinal cord collected from *ATF6 fl/fl; PLP-Cre-* and *ATF6 fl/fl; PLP-Cre+ mice.* Representative images of oligodendrocyte marker TPPP and ATF6 immunostaining in lumbar spinal cords of *ATF6 fl/fl; PLP-Cre-* mice (A,B) and *ATF6 fl/fl; PLP-Cre+ mice* (C,D).

| Cell Marker | Color | Clone | Vendor |
|---|---|---|---|
| **MHCII** | PE | M5/114.15.2 | Thermo Fisher |
| **Foxp3** | PE | FJK-16S | Invitrogen |
| **CCL3** | PE | DNT3CC | Thermo Fisher |
| **CD5** | PE | 53-7.3 | Thermo Fisher |
| **CD40** | PE-Cy7 | 3/23 | BioLegend |
| **IL-10** | PE-Cy7 | JES5-16E3 | BioLegend |
| **IL-17** | PE-Cy7 | 17B7 | Thermo Fisher |
| **Sca-1** | PE-Cy7 | D7 | Thermo Fisher |
| **CD45** | PE-C5 | 30-F11 | Thermo Fisher |
| **F4/80** | FITC | BM8 | BD |
| **VLA-4** | FITC | R1-2 | Thermo Fisher |
| **CD44** | FITC | IM7 | BD |
| **CD19** | FITC | 1D3 | Thermo Fisher |
| **Ly-6C** | PerCP-Cy5.5 | AL-21 | BD |
| **Lag-3 (CD223)** | PerCP-Cy5.5 | C9B7W | BD |
| **IFNg** | PerCP-Cy5.5 | XMG1 .2 | Thermo Fisher |
| **CD1d** | PerCP-Cy5.5 | 1B1 | BioLegend |
| **CD86** | BV421 | GL-1 | BioLegend |
| **CD80** | eFluor450 | 16-10A1 | Thermo Fisher |
| **CTLA-4 (CD152)** | PB | UC10-4B9 | BioLegend |
| **VID stain** | BV510 | L34966 (Cat #) | Thermo Fisher |
| **CD8** | BV605 | 53-6.7 | BD |
| **Ki67** | BV605 | 16A8 | BioLegend |
| **PD-L1** | BV650 | MIH5 | BD |
| **CD4** | BV650 | RM4-5 | BD |
| **CD11c** | BV711 | HL3 | BD |
| **CD3** | BV711 | 145-2C11 | BD |
| **B220** | BV711 | RA3-6B2 | BD |
| **CD11b** | BV786 | M1/70 | BD |
| **CD25** | BV786 | PC61 | BD |
| **Ly-6G** | PE-CF594 | 1A8 | BD |
| **PD-1 (CD279)** | PE-CF594 | J43 | BD |

**Table S1: Cell markers used in flow cytometry.**

| Target gene | Direction | Primer sequence (5' to 3') |
|---|---|---|
| *Arg-1* | Forward | CTCCAAGCCAAAGTCCTTAGAG |
| | Reverse | AGGAGCTGTCATTAGGGACATC |
| *IL-10* | Forward | AAGGCAGTGGAGCAGGTGAA |
| | Reverse | ACAATCAGAATTGCCA TTGCAC |
| *IL-4* | Forward | ACAGGAGAAGGGACGCCAT |
| | Reverse | GAAGCCCTACAGACGAGCTCA |
| *IL-17* | Forward | ATGCTGTTGCTGCTGCTGAG |
| | Reverse | TTTGGACACGCTGAGCTTTGAG |
| *iNOS* | Forward | GCTGGGCTGTACAAACCTTCC |
| | Reverse | TTGAGGTCTAAAGGCTCCGG |
| *TNF-α* | Forward | GGCAGGTTCTGTCCCTTTCA |
| | Reverse | ACCGCCTGGAGTTCTGGAA |
| *IFN-γ* | Forward | TGAACGCTACACACTGCA TCTTGG |
| | Reverse | CGACTCCTTTTCCGCTTCCTGAG |
| *Grp78* | Forward | CACGTCCAACCCCGAGAA |
| | Reverse | ATTCCAAGTGCGTCCGATG |
| *Grp94* | Forward | TCGTCAGAGCTGATGATGAAGT |
| | Reverse | GCGTTTAACCCATCCAACTGAAT |
| *Pdia6* | Forward | TGCCACCATGAATCAGGTTCT |
| | Reverse | TCGTCCGACCACCATCATAGT |
| *Aft4* | Forward | TGGATGATGGCTTGGCCAGTG |
| | Reverse | GAGCTCATCTGGCATGGTTTC |
| *Erdj4* | Forward | TTAGAAATGGCTACTCCACAGTCA |
| | Reverse | TTGTCCTGAACAATCAGTGTATGTAG |
| *Nrf2* | Forward | CAGGCCCAGTCCCTCAATAG |
| | Reverse | TCAGCCAGCTGCTTGTTTTC |
| *HO-1* | Forward | GATAGAGCGCAACAAGCAGAA |
| | Reverse | CAGTGAGGCCCATACCAGAAG |
| *Nqo1* | Forward | TCTCTGGCCGATTCAGAGTG |
| | Reverse | CTCCCAGACGGTTTCCAGAC |
| *AMDA10* | Forward | CAGCATCTGACCCTAAACCAAAC |
| | Reverse | CAGATAGAACCTGCACATTGCC |

**Table S2: Primer sets use for qRT-PCR.**
